# Extensive horizontal transfer of transposable elements shape fungal mobilomes

**DOI:** 10.1101/2025.06.16.659975

**Authors:** Josje Romeijn, Iñigo Bañales Belaunde, Michael F. Seidl

## Abstract

Transposons impact eukaryotic genome size and evolution. Horizontal transfer of transposons (HTT) is important for their long-term persistence, but has only been systematically studied in animals, and thus the abundance, impact, and factors that shape HTTs in lineages outside animals is unknown. Fungi are at least as ancient and diverse as animals and are characterized by extensive genome size variation caused by transposons. Here, we screened 1,348 genomes across fungal biodiversity, genome sizes, and lifestyles to detect extensive HTTs, that generated on average 7% but up to 70% of the transposon content in some taxa. We in total identified at least 5,518 independent HTTs, mostly involving *Tc1/Mariner* DNA transposons. While the majority of HTTs occur between closely related taxa, irrespective of their lifestyles, HTTs were particularly common in Mucoromycotina, Sordariomycetes, Dothideomycetes, and Leotiomycetes. Importantly, species lacking fungal-specific defense mechanisms against transposons and those with gene-sparse and repeat-rich genomic compartments are involved in significantly higher number of HTTs, unveiling ecological and genomic factors shaping HTTs. Our findings thus illuminate the dynamic landscape of HTTs in fungi, providing the framework to further study the impact of HTTs on genome evolution and the processes that mediate transposon transfers within and between eukaryotic lineages.

## Introduction

Transposable elements (TEs) are mobile DNA sequences that are highly abundant in eukaryotes, and in some cases comprise over 90% of their genomes^1,2^. Eukaryotes differ considerably in their mobilomes – the diversity and abundance of all TEs in their genomes. TEs have long been thought to be merely parasitic genomic sequences^3,4^, yet they are now seen as integral for genome evolution, shaping genome structure and content^5,6^ as well as genome functioning^7–9^. If left unchecked, TEs are able to rapidly increase in copy number^10^, and this proliferation is the main source of genome size expansions and variation in eukaryotes^11–13^.

To suppress TE activity, eukaryotes have evolved genome defense mechanisms, such as DNA silencing^14–16^. Since silenced DNA has a higher mutation rate due to less efficient DNA repair processes^17^, this DNA is prone to degeneration^18^. TE defense mechanisms are generally effective, as roughly 95% of TEs in eukaryotic genomes are fragmented^19^. Beyond host genome defenses, TEs are also purged due to their own transposition dynamics^20^, through hyperparasitism of non-autonomous TEs^21^. Moreover, since TE activity has often been considered to decrease host fitness, TEs are frequently eliminated due to selection or drift^22^. With mostly TE fragments being retained in eukaryotic genomes^19^, the evolutionary dynamics of eukaryotic mobilomes and processes contributing to their persistence remain enigmatic.

The evolutionary dynamics of eukaryotic mobilomes are traditionally viewed through the lens of Darwinian evolution by vertical descent and genetic conflicts between selfish TEs and the host genome^23,24^. However, the persistence of TEs over longer timescales implies that they can evade their seemingly inevitable extinction resulting from inactivation or elimination. The observation that some TEs such as the *P* element in flies could horizontally transfer to new hosts provides an intriguing mechanism to explain the long-term persistence and propagation of TEs^23,25,26^. Systematic quantification of horizontal transfer of transposable elements (HTTs) in insects uncovered that HTT-derived TEs account for up to 24% of the genome^27^, highlighting the substantial impact of HTT on insect mobilomes. While incidental HTT events have also been reported in animals, plants, and fungi^28–34^, systematic studies have been limited to few animal lineages, especially to insects and vertebrates^27,35^, with a more recent studies extending to annelids, arthropods, mollusks, chordates^36^, and flies^37^.

Fungi are well known ecosystem architects, pathogens of plants and animals, and cell factories in the bio-industry^38–42^. They emerged between 700 million and 1 billion years ago and represent one of the oldest and most diverse eukaryotic lineages. Their mobilomes are highly dynamic and are thought to contribute to their remarkable capacity for adaptation to a variety of environmental niches^43–47^. Notably, fungi lack protective germlines and, even though non-self-recognition systems exist^48^, cells of different fungal species can fuse^49–54^, potentially making them receptive to HTTs. Giant mobile genetic elements, called Starships, have been recently identified, and these TEs can transfer traits, incl. pathogenicity, between fungal lineagues^30,55,56^. Despite their significance^57^, they only constitute a minority of the mobilome and are only present in a subset of fungal lineages. To our knowledge, the abundance and impact of HTTs in fungi has not yet been systematically studied^58^. Here, we query the mobilomes of 1,348 fungal genomes, covering the entire kingdom to identify extensive HTTs and to uncover ecological, taxonomic or genomic factors that shape HTTs and fungal mobilomes.

## Results

### Independent transposable element-mediated genome expansions in the fungal tree of life

The sizes of fungal genomes and mobilomes differ by multiple magnitudes^59–62^, and we hypothesized that this variation is at least partially caused by HTTs. To study mobilomes and HTTs, we first obtained 1,348 publicly available fungal genome assemblies (**Tab. S1**), covering taxonomic and lifestyle diversity (**Fig. 1a**, **Tab. S2**), and captured the individual mobilomes using a combination of *de novo* and homology-based TE detection and annotation (**Tab. S3, S4**). On average, fungal genomes are 45.9 Mb in size, of which 11.8% are TEs (**Fig. 1a, Tab. S1**). Genome and mobilome size are correlated (Spearman’s rho = 0.65, p<1e-16, **Fig 1b**), corroborating that TEs are the drivers of genome expansions; genomes and mobilomes exhibit 100-fold size variation, e.g., in Pucciniomycotina^63,64^, Zoopagomycota^65^, or Dothideomycetes^66,67^. Notably, the ten largest genomes (>200 Mb) occur in multiple lineages and are characterized by large mobilomes (between 76 Mb to 1.1 Gb) (**Fig. 1b**), indicating that TE-driven genome expansions occurred independently across the fungal tree of life.

**Figure 1.**
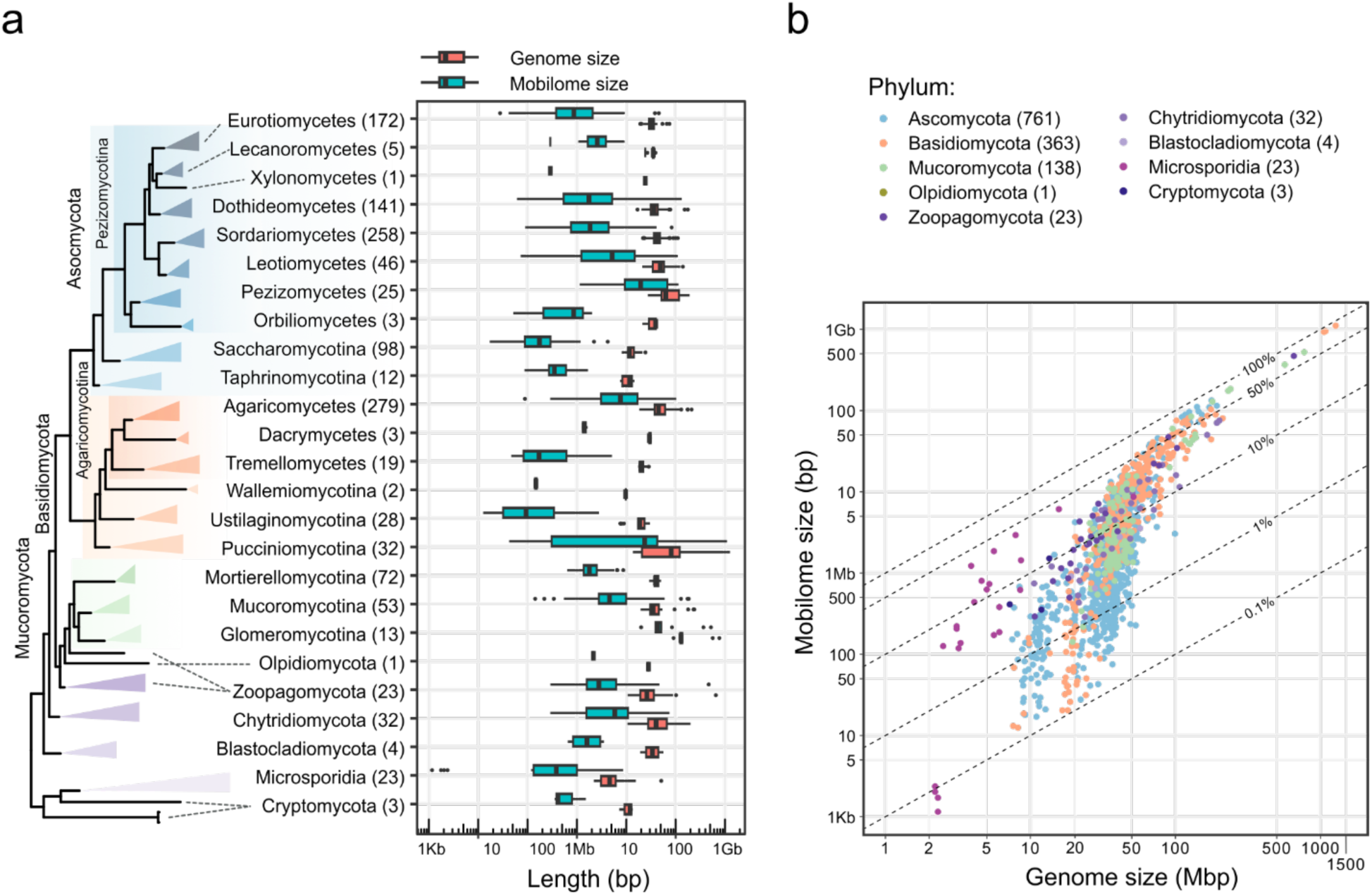
Transposable element-mediated genome expansions in the fungal tree of life. **a**, The 1,348 selected fungal genome assemblies (Tab. S1) cover the previously described fungal taxa, Cryptomycota, Microsporidia, Blastocladiomycota, Chytridiomycota, Zoopagomycota, Mucoromycotina, Basidiomycota, and Ascomycota^68^. The topology of the fungal tree of life was determined by a maximum-likelihood phylogeny based on 151 single-copy orthologous genes rooted between Microsporidia and Cryptomycota as well as the other phyla; the numbers in brackets indicate the number of genomes considered for each taxonomic lineage. Genome (red, in bp) and mobilome size (blue, in bp covered by TEs in a genome), vary up to 100-fold, ranging from 2.2 kb to 1.3 Gb and 1.1 kb to 1.1 Gb, respectively, with extensive variation within different fungal lineages. **b,** Genome and mobilome sizes are positively correlated (Spearman’s rho = 0.65, p<1e-16), consistent with TEs being the major driver for fungal genome expansions. Dashed lines indicate the proportion of fungal genomes being covered by TEs.

### Extensive horizontal transfer of transposable elements in fungi

To determine how HTTs contribute to fungal mobilomes (**Fig. 1**), we developed a computational approach (**Fig. S1**), inspired by recent work on insects and vertebrates^27,35^. We successfully detected HTTs linking nearly 65,000 TEs, representing 3.4% of all TEs with TE-related protein domains and >300 bp in length (*n* = 1,905,761, **Fig. S1a-c**; **Tab. S5**). By contrasting divergence, approximated by synonymous substitution rate (*Ks*), between TEs and single-copy orthologous genes, which are assumed to be inherited vertically, we detected in total ∼1.3 million candidate HTTs in around half (663) of all genomes. Since multiple candidate HTTs could be caused by the same event, we clustered those into communities (**Fig. S1d**), yielding 11,001 hit communities (**Fig. 3**; 2-3,908 TEs per community). To detect whether HTTs occurred in the common ancestor of taxa, we connected communities (**Fig S1e**) and subsequently compared those clusters to the species tree to determine the minimum number of HTTs (**Fig. S1e,f**). In total, we detected at least 5,518 independent HTTs, highlighting that HTTs occurs frequently between fungi.

**Figure 2.**
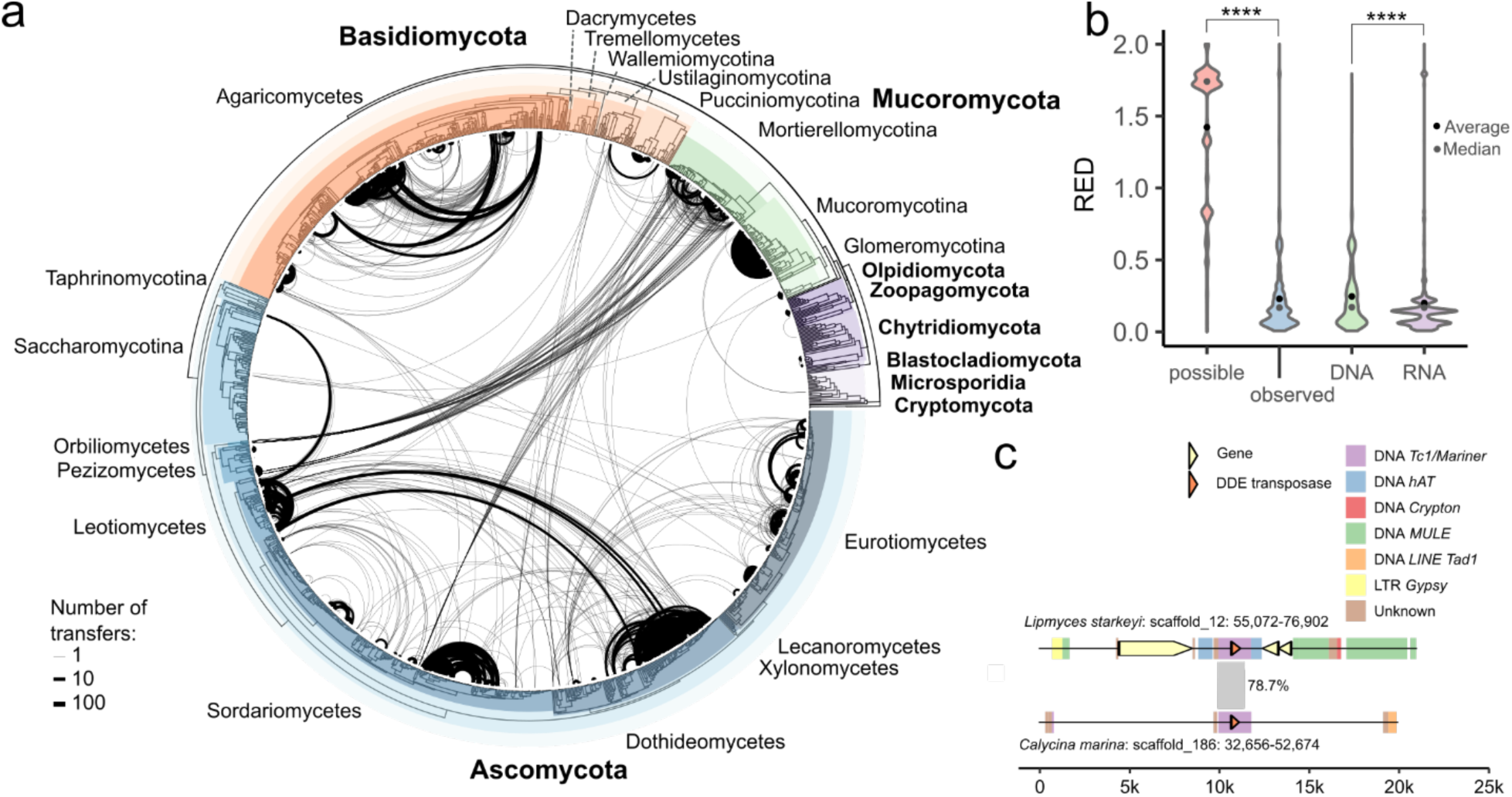
Horizontal transfer of transposable elements is widespread among fungi. **a**, HTTs occur throughout the fungal kingdom. HTTs between taxa (*n* = 11,001) in the fungal tree of life (for details, see Fig. 1a) are shown as links between taxa, the line width indicates the number of transfers. Extremely recently diverged taxa were treated as a single entity, and transfers between those were projected onto a single member. Using PhyloRank, relative evolutionary divergence (RED) values were calculated so that branches reflect relative time rather than substitutions. **b**, HTTs occurs more frequently between closely related taxa, particularly for RNA transposons. Distribution of RED values between genome comparisons between all possible genome comparisons are indicated in the boxplot, treating recently diverged taxa as a single entity, (possible, *n* = 916,245), observed genome comparisons involved in HTTs (observed, *n* = 11,001), HTTs of DNA transposon (DNA, *n* = 6,699), and HTTs of RNA retrotransposon (RNA, *n* = 3,874). RED values of 0 indicate close phylogenetic relationships and values of 2 indicate very distant phylogenetic relationships. Black dots and grey dots refer to the average and median, respectively. RED values of observed transfers are significantly lower than theoretically possible RED values, and RED values of RNA transfers compared are significantly lower than those for DNA transfers (one-sided Wilcoxon rank sum test, p < 1e-308 and p = 7.8e-64, respectively). **c**, Alignment of a candidate HTT involving two DNA *Tc1/Mariner* transposons located in *Calycina marina* and *Lipomyces starkeyi*, belonging to Leotiomycetes and Saccharomycotina and diverged roughly 500MYA^69^. Genome alignments of the genomic regions, inc. 10 kb up and downstream of TEs, revealed that the only alignment involved the HTT-derived TEs (78.7% identity).

**Figure 3.**
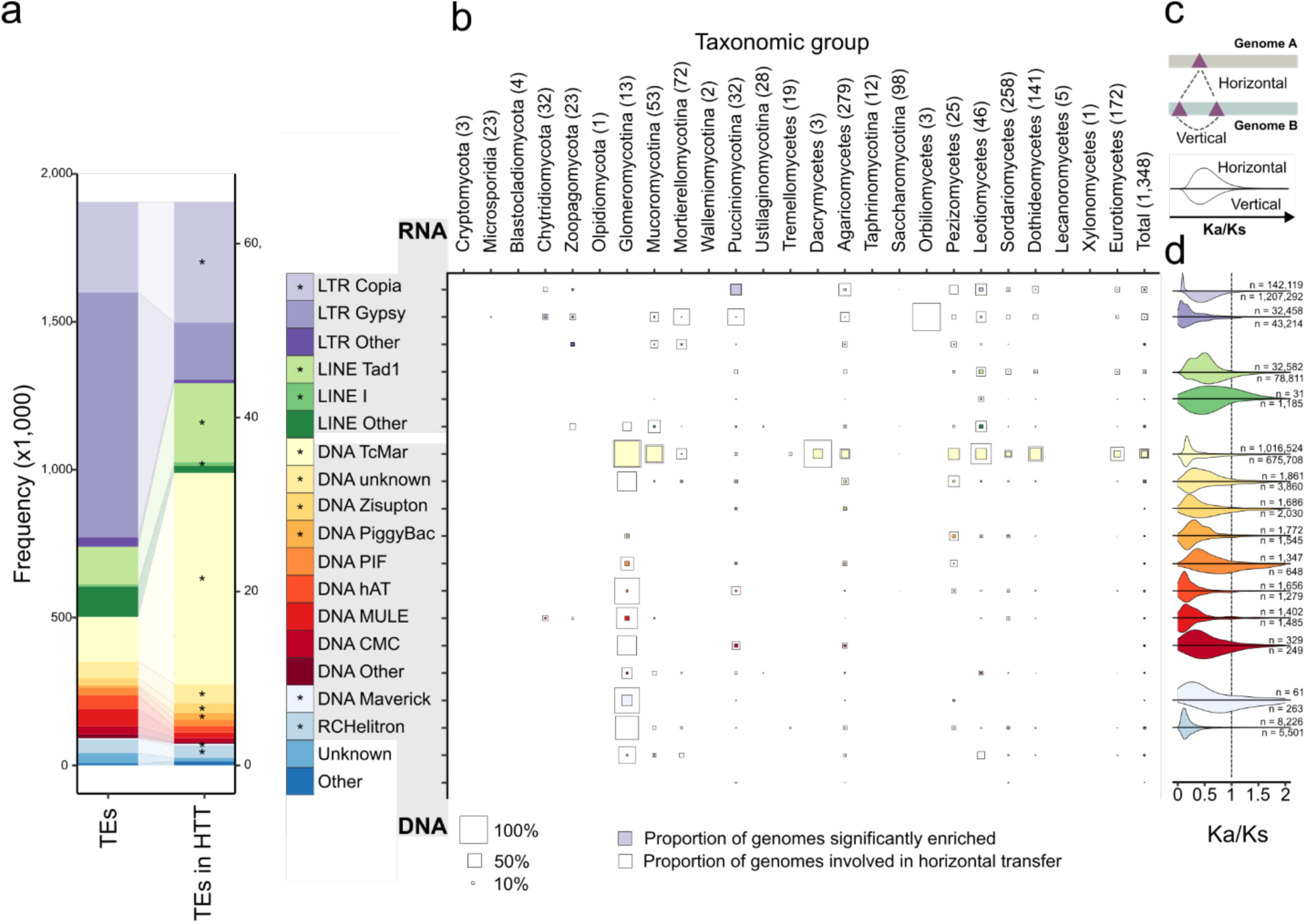
DNA *TcMariner*, LINE *Tad1*, and LTR *Copia*, are abundantly horizontally transferred and evolve under strict selective constraint, irrespective of their mode of transposition. **a**, Several TE superfamilies are enriched in TE-involved HTT. The number of transposable elements (TEs; TEs considered in the detection of horizontally transferred transposable elements (HTTs) encoding a TE-related protein domain and >300bp in length, *n* = 1,905,761, **Fig. S1**) as well as the number of TEs involved in HTTs (TEs in HTT; n = 64,815), separated by different TE classes and superfamilies. Significant enrichment of HTTs in a specific TE class/superfamily, determined by one-sided Fisher’s exact tests (**Table S4**), is indicated with an asterisk. **b,** DNA *Tc1/Mariner* is consistently enriched in individual fungal genomes in TEs involved in transfer (involved in *Tc1/Mariner* HTT: *n* = 447 genomes; enriched for TEs involved in *Tc1/Mariner* HTT: *n* = 300 genomes). Tile plot showing the occurrence of horizontal transfers in proportion of genomes in taxonomic groups per TE type (uncolored, “outer” squares), as well as the proportion of genomes in taxonomic groups enriched for TEs involved in transfer per TE type (colored, “inner” squares). Sizes of the squares indicate proportion size. The final row indicates the proportions of horizontal transfer and enrichment of TEs involved in transfer per specific TE type throughout all analyzed genomes. **c,** Cartoon explaining the differences between horizontally transferred TEs (∼65,000 TEs involved in ∼1.3 million candidate HTTs) and vertically transmitted TEs (based on TEs located in same genomes and present in same hit community). **d**, Fungal TEs are consistently under stringent purifying selection, irrespective of their mode of transposition. Evaluation of selection regimes acting on TEs, inferred through calculation of Ka/Ks values of TE-related protein domains located on horizontally and vertically transmitted TE pairs. Number of TE pairs per TE type are noted for horizontally and vertically transmitted TE pairs. TE pairs sharing <100AA of TE-related protein domains were excluded.

To investigate the HTT distribution, we projected ∼11k HTTs between extant species pairs (i.e., hit communities) onto the fungal phylogeny (**Fig. 2a**). HTTs are widespread (**Fig. 2a**), and occur significantly more frequent between closely related lineages (one-sided Wilcoxon rank sum test, p < 1e-308 and p = 7.8e-64, respectively, **Fig 2b**), a pattern similarly observed in animals^27,35,36^. While nearly all HTTs (10,785 out of 11,001 HTTs) occur within the same taxonomic group (**Fig. 4b**). For example, we observed the transfer of a *DNA Tc1/Mariner* transposons between *Calycina marina* and *Lipomyces starkeyi*, belonging to Leotiomycetes and Saccharomycotina that diverged roughly 500 MYA^69^. We even observed 99 transfers between taxa that diverged >700 MYA^69^ (**Fig. 2c, 4b**), emphasizing that HTTs can even occur between highly divergent taxonomic groups.

**Figure 4.**
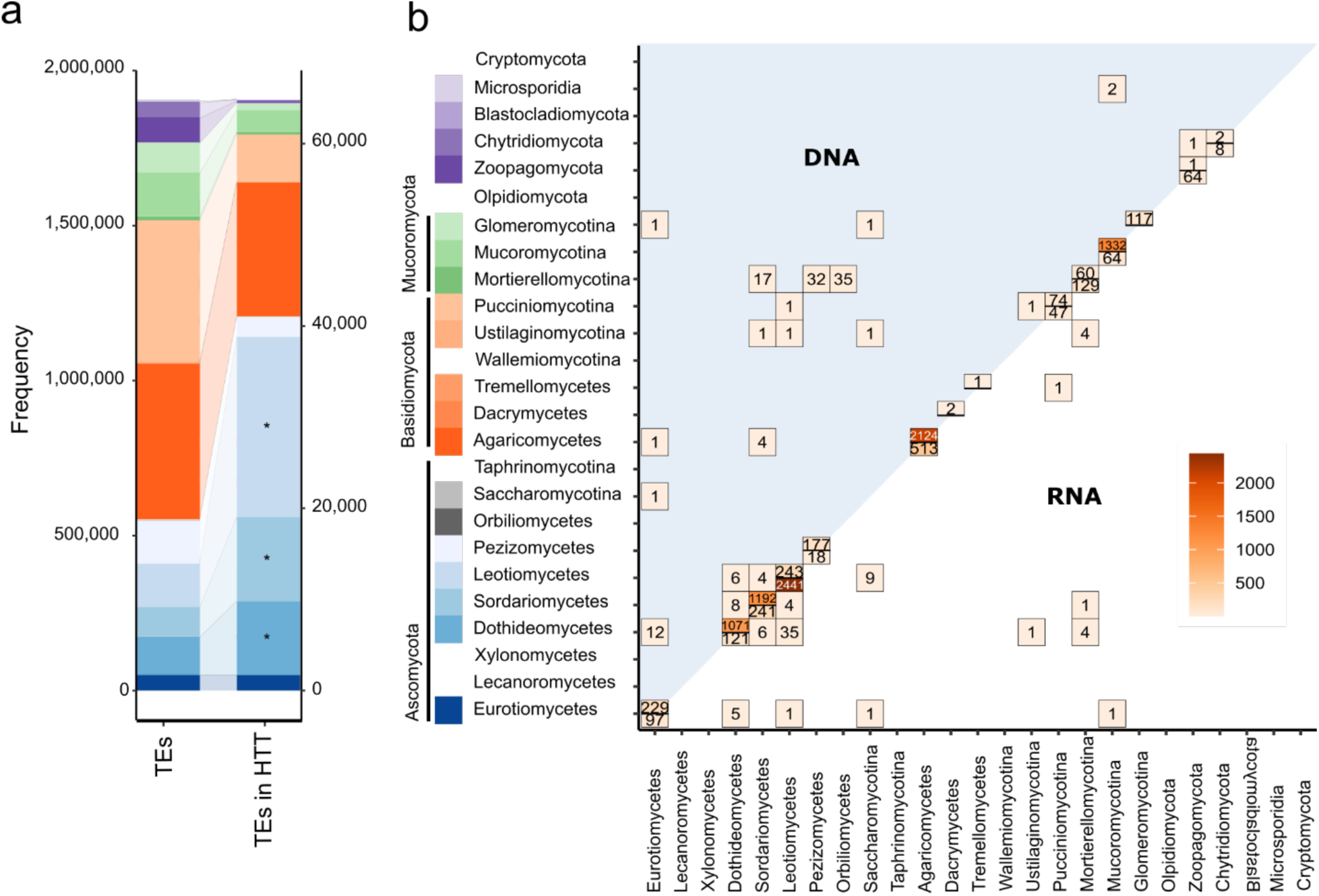
Horizontal transfers of transposable elements are enriched in few taxonomic lineages. **a**, Sordariomycetes, Leotiomycetes, and Dothideomycetes are enriched in TEs involved in horizontal transfer. The number of transposable elements (TEs; TEs considered in the detection of horizontal transfer of transposable elements (HTTs), *n* = 1,905,761, **Fig. S1**) as well as the number of TEs involved in HTTs (*n* = 64,815) are shown, separated by different fungal lineages. Significant enrichment of HTTs in a specific fungal lineage, determined by one-sided Fisher’s exact tests (**Tab. S8**), is indicated with an asterisk. **b,** HTTs occurs more often between closely related lineages, and each lineage favors transfer of either DNA or RNA retrotransposons. The number of HTTs between different taxonomic lineages is visualized as the number of hit communities. The upper and lower triangle shows transfers of DNA transposons and RNA retrotransposons between fungal lineages, respectively.

### *Tc1/Mariner* are abundantly horizontally transferred transposable elements in fungi

TEs are broadly classified into retrotransposons and in DNA transposons^70^. We observed that both types are involved in HTTs in fungi, similar to vertebrates and insects^27,35,36^. While retrotransposons are more abundant in fungi, more DNA transposons have been transferred (**Fig. 2b,3a**). Several TE orders/superfamilies such as the DNA *Tc1/Mariner*, LTR *Copia*, or LINE *Tad1* are enriched in HTTs (one-sided Fisher’s exact test, p < 1e-308, p = 2.3e-287 and p < 1e-308, respectively; **Fig. 3a**, **Tab. S7**). Frequent HTT of *Tc1/Mariner* and *Copia* have also been observed in animals^27,35,36^, suggesting a shared pattern across eukaryotes. However, LINE *Tad1* is only frequently involved in HTTs in fungi. Interestingly, *Tc1/Mariner* were involved in HTTs in 447 genomes (33% of all genomes), spanning nearly all lineages, *Tc1/Mariner* were also significantly enriched in HTTs in two third of these (300 genomes) (**Fig. 3b**, **Fig. S2**). By contrast, LTR *Copia* and *Gypsy* are highly abundant in fungi (**Fig. 3a**), and were also found to be transferred in 272 and 285 genomes (>18% of all genomes), respectively, but were only enriched in less than 5% of all genomes (**Fig. S2**), suggesting that *Tc1/Mariners* are more frequently involved in HTTs or that these are much more successful at proliferation after a HTT compared to other, more abundant TEs.

### Transposable elements, irrespective of their mode of transposition, evolve under purifying selection

TEs might evolve under different selective constraints depending on their mode of transposition (vertical vs. horizontal)^35^; horizontally transferred TEs need to encode proteins essential for successful transposition in the absence of those in the recipient. We estimated the selection regimes using the ratio non-synonymous mutation rates (*Ka*) to synonymous mutation rates (*Ks*) (**Fig. S1**) for 1.3 million horizontally and 2.1 million vertically transmitted TEs (**Fig. 3c**). Apart from vertically transmitted *LINE I* and horizontally transmitted DNA *PIF-Harbinger* and *Mavericks*, all TEs experience purifying selection (one-sided Wilcoxon rank sum test, **Tab. S9**), irrespective of their mode of transmission. However, comparison between horizontally and vertically transmitted TEs revealed that horizontally transferred TEs are subject to stronger purifying selection across all RNA retrotransposons, except *LINE I*, and in DNA transposons *Tc1/Mariner*, *Zisupton*, *PIF-Harbinger*, *Maverick,* and *RC Helitron* (one-sided Wilcoxon rank sum test, see **Tab. S10**). In contrast to vertically transmitted *Tc1/Mariner* and *hAT* in vertebrates^35^, those TEs evolve under purifying selection in fungi (Ka/Ks < 0.65, **Tab. S9**). The high abundance of DNA transposons in vertebrates might allow for *trans*-preference as TEs with non-functional transposases could rely on transposases of other copies^21^. The lower prevalence of DNA transposons in fungi (**Fig. 3a**) could limit transposase availability, explaining the stronger purifying selection on those TEs.

### Horizontal transfers of transposable elements are enriched in few taxonomic lineages

HTTs are widespread in fungi (**Fig. 2a**), but some lineages might be more frequently involved in HTTs. To test this, we performed enrichment analyses comparing TEs involved to those not involved in HTTs (**Fig. 4a**). Interestingly, mobilomes of Dothideomycetes, Sordariomycetes, and Leotiomycetes are significantly more often involved in HTTs than other taxonomic groups (**Fig. 4a**). However, this does not distinguish whether these lineages are more frequently involved in HTTs, or whether transferred TEs are simply more successful at proliferation after the initial transfer. To disentangle these two, we compared the number of HTTs each genome participates in, measured as the number of hit communities. This revealed that Dothideomycetes have significantly more transfers per genome compared to Eurotiomycetes (one-sided pairwise Wilcoxon rank sum tests, p = 7.7e-4, **Fig. S3**) but not compared to other fungal lineages. Leotiomycetes and Sordariomycetes do not differ significantly from other lineages (**Fig. S3**), suggesting that while HTTs are not more frequent in these groups, transferred TEs can still successfully proliferate, also seen, though less strongly, in Dothideomycetes. Mucoromycotina, in contrast, are not enriched in TEs involved in HTTs, but genomes participate in significantly more transfers compared to other lineages (**Fig. 4a**, **Fig. S3**), implying that transferred TEs do not increase in abundance (**Fig. 4a**). Other lineages with large mobilomes, e.g., Agaricomycetes or Pucciniomycotina (**Fig. 1a**), are not enriched in HTTs and do not have abundant HTTs per genome (**Fig. 4a, Fig. S3**), highlighting that large mobilomes do not imply more HTTs or successful proliferation of TEs after transfers.

We then determined if specific TE classes transfer preferentially between some lineages, as the vast majority of all HTTs occur within the same lineage **(Fig. 2b,4b**). Remarkably, Mucoromycotina, Agaricomycetes, Sordariomycetes, and Dothideomycetes predominantly transfer DNA elements (5,719 out of 6,618), while Leotiomycetes nearly exclusively transfer retrotransposons (2,441 out of 2,684), indicating that lineages are biased towards transfer of either DNA- or retrotransposons. Notably, 8,951 out of 11,001 HTTs occur between taxa of the same lifestyles (**Fig. S4**), suggesting that the ecological niche in which species reside plays a role in shaping HTTs.

### The absence of genome defenses is correlated with an increase in horizontally transferred transposable elements

Even though common in fungi (**Fig. 2a**), HTTs are only enriched in few taxa (**Fig. 4**), suggesting intrinsic differences that impact HTT frequency or the success of HTT-derived TEs. Genome defenses, such as DNA silencing via histone or DNA modifications as well as the fungal-specific repeat-induced point (RIP) mutation pathway^71,72^ could impact HTTs. We classified genomes as either RIP-proficient or RIP-deficient (**Fig. 5a**), which demonstrates that most Ascomycetes are, as expected^73^, RIP-proficient; the presence of five key enzymes involved in TE defense mechanisms, either in DNA or histone methylation as well as in RIP are correlated with higher proportions of RIP-affected repeats^74–79^ (one-sided Wilcoxon rank sum test, see **Fig. S5**). Across fungi, we observed a significant enrichment of the number of TEs involved in HTT in RIP-proficient genomes (one-sided Fisher’s exact test, p = 1.7e-181, see **Fig. S6**), which is counter intuitive as RIP-proficient genomes are expected to be efficient in eliminating TEs. Most likely this is a taxonomic signal, as RIP occurs mostly in Ascomycetes^73^ but only in few Basidiomycetes^80,81^, but nearly 40% of HTTs occur within Basidiomycetes. Interestingly, in Ascomycete lineages enriched in HTTs, i.e., Dothideomycetes, Sordariomycetes, and Leotiomycetes (**Fig. 4a**), RIP-deficient genomes are significantly enriched in HTTs as well as in HTT-derived TEs (one-sided Wilcoxon rank sum test, p = 0.0028, **Fig 5d**; one-sided Fisher’s exact test, p < 1e-308**, Fig. S6**), demonstrating that absence of RIP does not only correlate with reduced genome defenses against TE proliferation^82^ and activity^83^, but also impacts HTTs.

**Figure 5.**
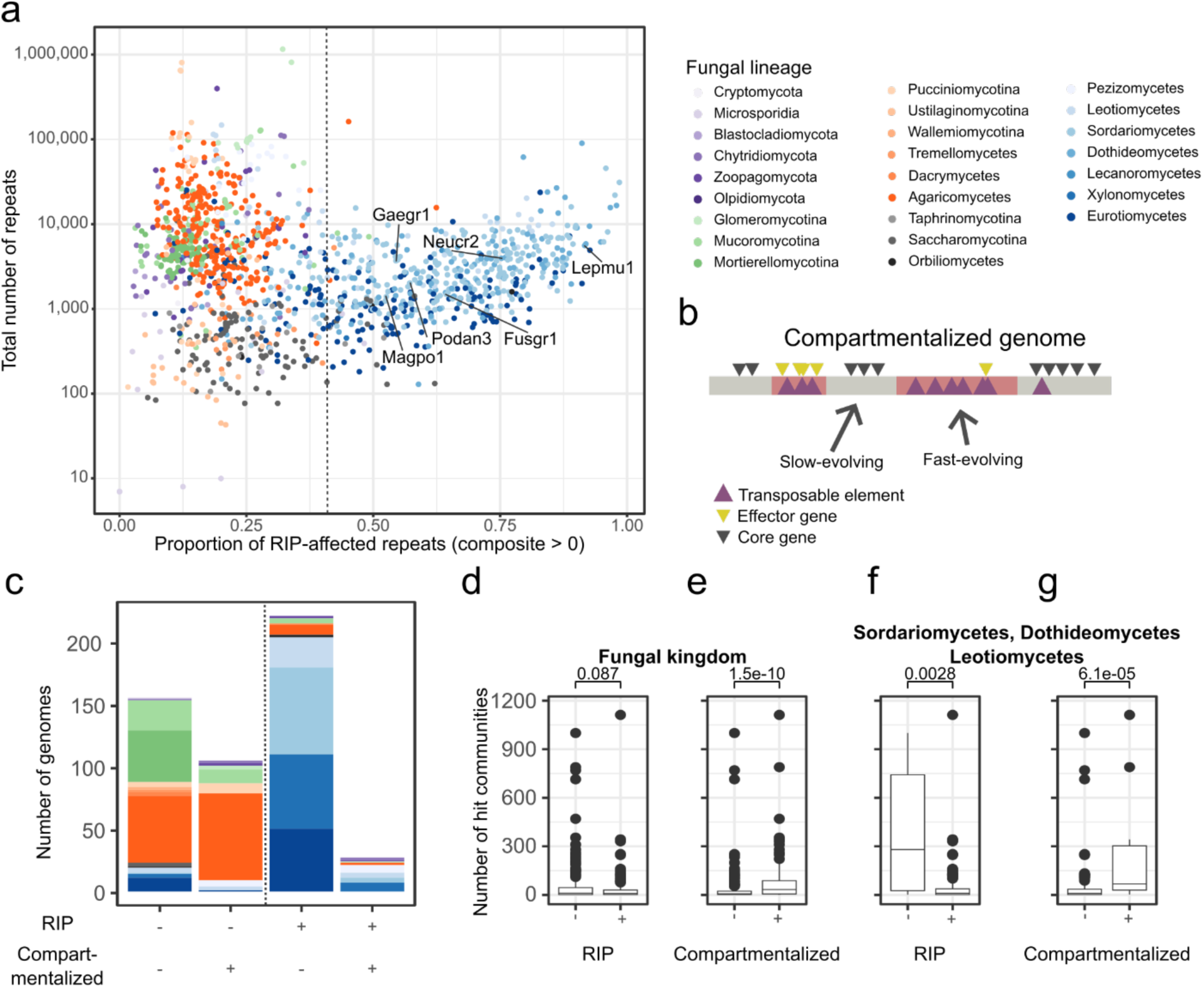
Genome compartmentalization and loss of Repeat induced point mutations (RIP) are associated with an increase in horizontal transfers of transposable elements (HTT). **a**, The RIP proficient taxa are almost exclusively found within Ascomycetes, as determined by the proportion of repeats that bear RIP-signatures. The total number of repeats (including TEs) per genome and proportion of TEs bearing signatures of RIP (composite index > 1) per genome is displayed, colored by fungal lineage. Pie charts indicate the distributions of RIP-proficient (proportion of TEs bearing signature of RIP > 40%) and RIP-deficient genomes, We highlight few species that have been experimentally proven to be RIP-proficient: Magpo1; *Magnaporthe oryzae*, Lepmu1; *Leptoshpaeria maculans*, Fusgr1; *Fusarium graminearum*, Podan3; *Podospora anserina*, Gaegr1; *Fusarium solani*, Neucr2; *Neurospora crassa*^84–89^. **b**, A cartoon explaining genome compartmentalization in fungal genomes. Grey compartments denote slow-evolving, gene-dense, and TE-sparse regions, red compartments are fast-evolving, TE-dense and gene-sparse regions^90–100^. Note that effector genes are often located in TE-dense and gene-sparse regions. **c**, Number of genomes classified as RIP-proficient and RIP-deficient, as well as compartmentalized or non-compartmentalized. The dotted line separates RIP-proficient from RIP-deficient categories. **d**, Number of HTTs genomes participate in, visualized for RIP-proficient and RIP-deficient fungal taxa, **f**, and RIP-proficient and RIP-deficient genomes belonging to Sordariomycetes, Dothideomycetes, and Leotiomycetes, showing that Ascomycete genomes that have lost RIP are significantly more involved in HTTs. **e**, Compartmentalized genomes are significantly more involved in transfers. Genomes are considered to be compartmentalized if 5% of genes have an intergenic distance of >5 kb and a TE closer than 1 kb. Boxplot shows the number of HTTs genomes participate in, visualized for compartmentalized and non-compartmentalized genomes, **g**, and compartmentalized and non-compartmentalized genomes of Sordariomycetes, Dothideomycetes, and Leotiomycetes. P-values of one-sided Wilcoxon rank sum tests are included above for each comparison.

### Compartmentalized genomes are enriched in horizontally derived transposons

Many fungal genomes are compartmentalized, separating gene-dense and repeat-sparse regions, encoding core genes, from gene-sparse and repeat-rich regions^90,91^, with the latter often being enriched in lineage-specific orphan or pathogenicity-related genes^90–100^. We observed a consistent and significant enrichment of HTTs (one-sided Wilcoxon rank sum test, p = 1.5e-10, **Fig. 5e,f)**, but not an enrichment of HTT-derived TEs in taxa harboring gene-sparse and repeat-rich regions (one-sided Fisher’s exact test, p = 1, **Fig. S6**;), supporting the hypothesis that such genome organization facilitates HTTs but does not promote subsequent TE amplification. However, we did not observe that HTT-derived TEs were significantly closer to effector or orphan genes than other genes (one-sided Wilcoxon rank sum tests, **Tab. S12**). Thus, while compartmentalized genomes are enriched in HTT, there is currently no evidence for rampant proliferation of HTT-derived TEs or for HTT-derived TEs localizing predominantly in TE-rich genomic compartments.

### Horizontally transferred transposable elements shape a significant proportion of fungal mobilomes

Lastly, we quantified the impact of HTTs on genome and mobilome sizes (**Fig. 1**), using all HTT-associated TEs and TE fragments (**Fig. 6a**). The proportion of fungal mobilome derived from HTTs ranges from 0.004% to up to 67.9%, averaging at 7.3%. Interestingly, for 22% of genomes involved in HTT (i.e., 150 genomes), we could trace more than 10% and up to 70% of the mobilomes back to HTTs (**Fig. 6b,c**), and the total mobilome size is positively correlated with abundance of HTT-associated TEs (Spearman’s rank correlation, p = 2.2e-76, rho = 0.636; **Fig. 6b**). Taking protein coding-genes and other genomic region into account, the proportion of genomes derived from HTTs ranges from 0.0007% up to 51.6%, with an average of 1.5%. While compartmentalized genomes harbor significantly more TEs and HTT-associated TEs (one-sided Wilcoxon rank sum test, p = 6.2e-79 and p = 3.9e-33, respectively, **Fig. 6b**), they do not differ in the proportion of mobilome explained by HTTs (Wilcoxon rank sum test, p = 0.316, **Fig. 6c**). Thus, independent of fungal genome compartmentalization, these results demonstrate that HTTs are important in shaping fungal mobilomes.

**Figure 6.**
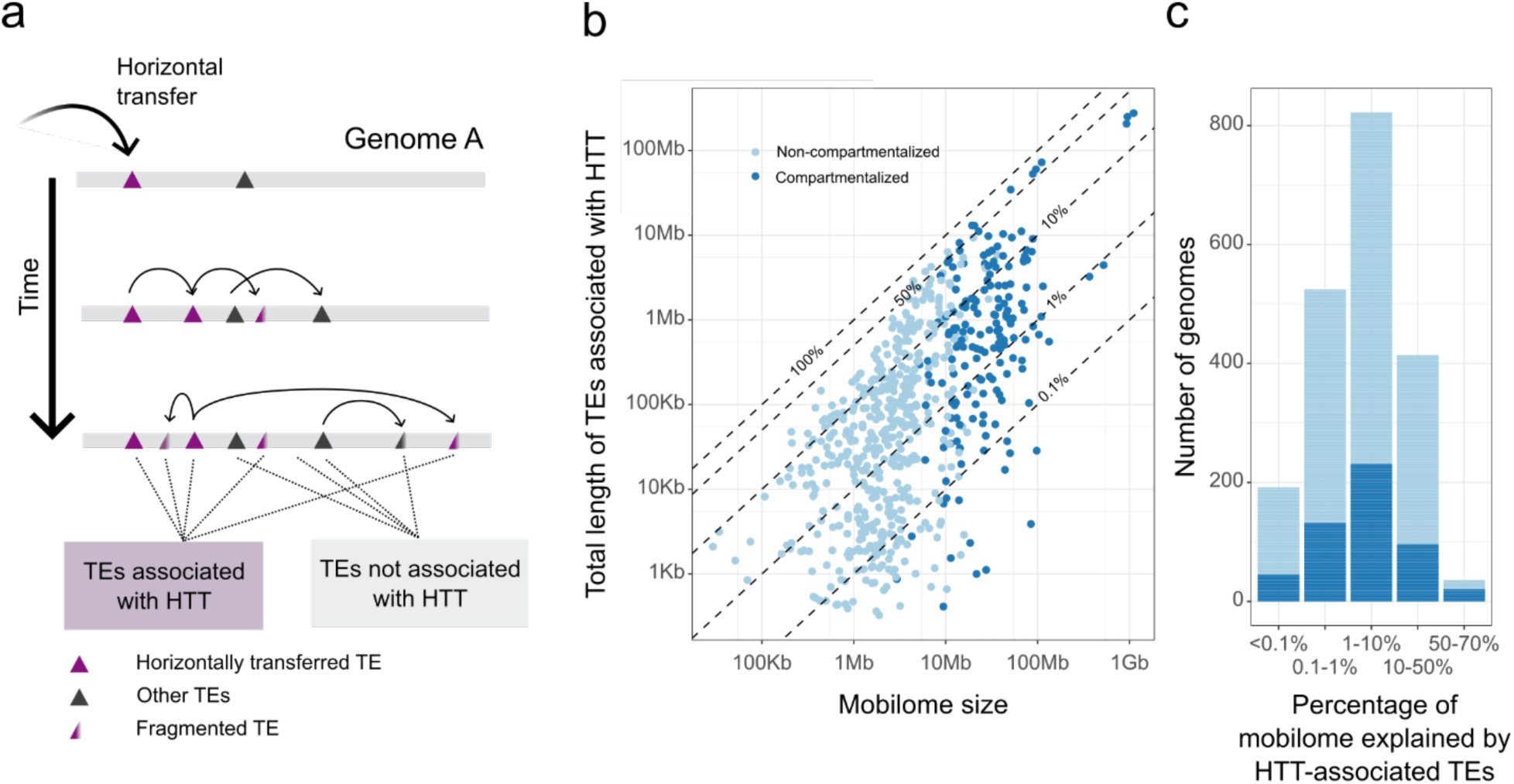
HTT-associated TEs explain up to 70% of fungal mobilomes. **a**, A cartoon illustrating how TEs can be associated with HTTs, including TE fragments, TEs not encoding TE-related protein domains, or TEs derived from original TE that had been involved in transfer. **f**, The total length of HTT-associated TEs is positively correlated with mobilome size (Spearman’s rank correlation, p = 2.2e-76, rho = 0.636), and compartmentalized genomes with repeat-rich, gene-sparse regions (indicated by colors) have significantly larger mobilomes and more HTT-associated TEs (one-sided Wilcoxon rank sum test, p = 6.2e-79 and p = 3.9e-33, respectively). **c**, Compartmentalized genomes do not differ in their proportion of mobilome explained by HTT compared to non-compartmentalized genomeso. Frequencies of proportions of mobilomes explained by HTT-associated TEs, colored by genome compartmentalization based on repeat-rich, gene-sparse regions.

## Discussion

Transposons are ubiquitous in eukaryotes, shaping their genome size and evolution^1,2,7,11,12,27,100^. Long-term persistence and proliferation is driven by their horizontal transfer^12,23,26^, evading their inevitable extinction resulting from genome defenses or selection. Horizontal transfer of transposable elements (HTT) has only recently been systematically studied in few animal lineages^27,35–37^, revealing that HTT is widespread and greatly impacts animal genomes^27,37^. However, it has not yet been studied systematically in other eukaryotic kingdoms^58^, raising important questions on the abundance and impact of HTTs as well as factors that shape HTTs in eukaryotes outside animals.

Here, using a conservative computational approach and a genomic dataset comprising 1,348 fungal genome assemblies spanning the breadth of the fungal kingdom, we demonstrate that HTT is widespread in fungi, occurring across nearly all studied phyla. HTTs in fungi had been previously reported incidentally such as for example the transfer of an RNA retrotransposon from fungi to plants^101^ or of an LTR element within the Saccharomycotina^102^. Recently, large cargo-carrying Starships have been discovered in fungi^55,56,103–105^, and recent experimental evidence suggest that they can undergo horizontal transfer between distantly related species belonging to the Eurotiomycetes^30^. We here focused on canonical DNA elements and retrotransposons and revealed that HTT is not a rare phenomenon but is rather widespread throughout the fungal kingdom.

We show that up to 70% of fungal mobilomes and 52% of their genome sequence can be explained by TEs associated with HTTs, emphasizing the pervasive influence of HTTs on fungi. In insects, up to 25% of their genomes is composed of TEs derived from HTTs ^27^, and up to 55% of the genomes of flies are composed of such TEs^37^, collectively suggesting that HTTs and their activities are pervasive in shaping eukaryotic mobilomes. High TE activity within host genome drives variation in genome size and content, and is often considered detrimental to the host’s fitness^26^. Therefore, eukaryotes evolved TE defense mechanisms, such as the fungal-specific RIP process, to suppress TE activities^14–16,71,72^. Loss of RIP in Leotiomycetes has been associated with an increase in genome size^82^. Our data extends this observation by linking the success of HTTs to the absence of RIP in Dothideomycetes, Leotiomycetes, and Sordariomycetes. However, RIP is also absent in lineages with many TEs, such as the Pucciniomycotina, or Chytridiomycota, that are not involved in abundant HTTs, suggesting that other mechanisms may restrict HTTs.

Many fungal genomes are compartmentalized, with repeat-rich and gene-sparse regions^90–100^. We observed that these genomic compartments are consistently associated with an increased number of HTTs, but not with abundant TE proliferation after HTT. Dynamic genome compartments, are often depleted of core genes, and are thought to serve as adaptive cradles subject to rapid genomic evolution^90,100^. These may act as permissive environments for TE activity, facilitating the integration of novel elements into the host genome. However, the functional consequences of HTT remain currently unclear as we do not find these HTT-derived TEs to be preferentially localized near orphan genes nor near genes involved in niche adaptation, such as those encoding secreted effector proteins^90–100^.

LTR *Copia* and LINE *Tad1* as well as *Tc1/Mariner* are transferred more frequently than expected given their overall abundance in fungal mobilomes. This pattern is consistent with prior observations, as many HTTs involving *Tc1/Mariner* have been also reported in animals^27,35,36^, and between prokaryotes and eukaryotes^106^, while LTR *Copia* transfers have been reported within plants^107,108^. HTTs of LINE *Tad1* seem to be unique to fungi, as these, to our knowledge, have not previously been reported. We note differences in evolutionary constraints acting on *Tc1/Mariner* across eukaryotic lineages, as these TEs appear to be subjected to much stronger purifying selection in fungi compared to vertebrates^35^. One possible explanation is the generally lower abundance of DNA transposons in fungi compared to vertebrate genomes^35^, thereby reducing the availability of transposases, resulting in stronger purifying selection to maintain functional transposases in DNA transposons.

HTTs occur across almost all fungal phyla, yet they are more frequent in Mucoromycotina, and proliferation of TEs subsequent to a HTT is more abundant in Dothideomycetes, Sordariomycetes, and Leotiomycetes. Furthermore, HTTs occur significantly more frequently between closely related species, a pattern also observed in animals and plants^27,35,36,109,110^. This ‘phylogenetic distance effect’ is not unique to intra-genomic parasites, such as TEs, as it is widely recognized in many host-parasite systems^111–114^. One possible explanation is that pathogens and parasites are more successful in (cellular) environments similar to their host, which are more likely to be found in closely related species^115^. Both parasitism and hyphal fusion, two mechanisms proposed to facilitate HTTs^23,50,58,116–118^, tend to occur more often between closely related species; parasites are generally host-specific^111,114^, and hyphal fusion is typically restricted to closely related species^49,51,53,118^. However, generalists can infect multiple, often diverse hosts^119^ and horizontal transfers of entire chromosomes between distantly related fungi pathogens have been experimentally demonstrated^120^. Moreover, hyphal fusion is thought to mediate the experimentally observed transfer of Starships^30^, contradicting the notion that such horizontal transfers are strictly limited to closely related taxa. Other proposed HTT vectors may further circumvent the phylogenetic distance barrier. For instance, viruses often infect multiple, distantly related hosts^121^. Some TEs, such as *Mavericks* or LTRs, are even suggested to be capable of forming capsid- or envelope-coated viral-like particles, potentially capable of surviving extracellular horizontal transmission^26,29,122–125^. Additionally, extracellular vesicles (EVs) are another intriguing HTT vector, and EV-mediated HTTs have been observed between bacterial genomes^126,127^. Some pathogenic fungi are capable of producing EVs that carry mRNA^128,129^, and cross-kingdom uptake of fungal EVs by plant and animal hosts, and *vice versa*, has been experimentally demonstrated^128–132^, suggesting that these processes could mediate HTTs in fungi as well as between fungi and other eukaryotic lineages. While our current study is limited to HTT detection within fungal kingdom, it remains an open and exciting question how frequently cross-kingdom HTTs occur and what impact these will have on eukaryotic genome evolution.

## Material and Methods

### Source data

To describe fungal mobilomes and systematically identify HTT, we obtained 1,348 publicly available genomes assemblies and their protein-coding gene annotation from the U.S. Department of Energy (DOE) Joint Genome Institute (JGI)^68^ (**Tab. S1)**, downloaded in August 2023. We applied NCBI taxonomy to determine fungal taxonomic lineagues^133,134^, and we used CATAstrophy^135^, which utilizes the carbohydrate-active enzyme (CAZyme) content, to predict trophic lifestyles for fungal genomes.

### Reconstruction of fungal tree of life

Core genes, i.e. single copy orthologous genes, were identified in fungal genomes assemblies using the BUSCO (Benchmarking Universal Single-Copy Orthologs) pipeline (v5.4.7)^136,137^, independently for 24 different fungal BUSCO datasets: one kingdom-level, four phylum-level, seven class-level, and twelve order-level, **Tab. S1**). A phylogenetic tree was constructed based 151 out of 758 core genes of the kingdom-level BUSCO set, using IQ-TREE (v2.2.6, settings: -B 1000 -m MFP -mset WAG+C60,LG+C60,Q.pfam+C60 -mfreq -mrate G4)^138^, allowing up to 10% missing genomes per core gene. We used FAMSA (v2.2.2)^139^ for construction of the amino acid alignment and trimAI (v1.4.rev15)^140^ to trim the alignment. The tree was constructed with 1,000 bootstrap replicates. ModelFinder was employed to identify the best substitution model^141^ from the following models: WAG+C60, LG+C60, or Q.pfam+C60, with G4 used for model selection. The tree was rooted between the Microsporidia + Cryptomycota on one side, and the other fungal taxa on the other side of the root. We used ggtree^142^, implemented in R (v 4.4.3), to visualize the phylogenetic tree.

To determine relative divergence times, we calculated the relative evolutionary divergences (RED) of genome assemblies^143^ in the tree using PhyloRank (v0.1.12) (https://github.com/dparks1134/PhyloRank), using the decorate function with option --skip_rd_ine and the outliers function.

### Detection and quantification of horizontally transferred transposable elements in fungi

The computational approach for HTT detection is inspired by the previous work on HTT identification in vertebrates and insects^27,35^. For computational efficiency, we used Snakemake (v8.12.0)^144^ to parallelize the computational pipeline.

### Synonymous mutation rate calculation of BUSCO genes

To estimate the evolutionary divergence between species, the synonymous mutation rate (*Ks*) of core genes was calculated for each node in the phylogenetic tree. First, the lowest taxonomic BUSCO set (see above) was determined for each node, and secondly, for each BUSCO gene, a representative core gene from each daughter clade was selected based on its length (or randomly, in case of equal length) using the ete3 package (v3.1.3)^145^, implemented in Python (v3.6.13). The two amino acid sequences of core genes were aligned using mafft (v7.505)^146^, and used to construct a codon alignment with PAL2NAL (v14, -nogap)^147^. Lastly, *Ks* values were estimated for each core gene with the *kaks* function in the seqinr (v4.2-36) R-package. Per node, the 0.5% quantile was estimated from the distributions of *Ks* values. To conservatively determine horizontal transfers, comparisons between species were only allowed in nodes with a 0.5% quantile larger than 0.01 of their respective BUSCO *Ks* distribution. Nodes failing to meet this criterion were subsequently collapsed, and species inside a collapsed node are treated as a single taxonomic entity. A total of 368 species were collapsed into 126 clades, with most collapsed clades comprising only two species (**Tab. S13**). We continued the analysis with 1,106 clades, including collapsed clades as well as genome assemblies not located in collapsed clades.

### TE annotation, classification, and identification of HTT candidates

TEs were annotated in fungal genomes using EarlGrey^148^, which included TE classification based on RepeatMasker^149^, complemented with TE classifications based on PASTEC (v2.0)^150^; RepeatMasker classifications were favored for conflicting classifications. We filtered out simple repeats, low complexity regions, and G/A/GA-rich regions. This yielded a total amount of 16,410,702 million TEs, which were subsequently extracted from genome assemblies with BEDtools (v2.31.1)^151^.

To accurately estimate evolutionary divergence, *Ks* rates were calculated over protein domains. We thus compiled a list of TE-related protein domains (**Tab. S14**), based on previous studies^152,153^. TEs with a length less than 300bp were discarded, after which Transeq (v6.6.0.0, -frame=6)^154^ was used to translate TE DNA sequences into all six open reading frames. Next, RPSTBLASTN (v2.12.0+, -evalue 0.001 - max_target_seqs 1)^155^ was used to search 142 Conserved Domain Database (CDD)^156^ domains against TE sequences with a cut-off e-value of 0.001, followed by HMMER (v3.4, -E 0.001) (hmmer.org) to search to 110 Pfam Hidden-Markov Model (HMM) profiles against the translated TE sequences (**Table S14**). Only TE copies containing a TE-related protein domain were kept for further analyses, resulting in 1,905,761 TE copies on 1,348 fungal genomes.

To decrease the computational load, identical TEs were identified per genome. Per identical set of TEs, only one representative sequence was kept for further analysis, resulting in 1,897,358 TEs. Subsequently, we performed an all-vs-all BLASTn (2.12.0+, -dbsize 10,000,000,000 -evalue 10 -max_target_seqs 100,000)^155^ search of TE sequences. Only hits between clades with a percent identity of >75%, a length of >300bp, and a bitscore of 200 were kept, resulting in a total of 194,575,775 blast hits. Subsequently, fragmented blast hits were chained with a maximum gap and maximum overlap of 600 bp, and to avoid perfect but partial hits, only blast hits were kept for which the alignment covered both TEs by 60%. After ensuring the blast hits included only one-directional hits, 14,591,977 blast hits remained for further analysis.

*Ks* was estimated for each blast hit, using the same methods described above for BUSCO genes. Protein domains had to be present in both TEs at precisely the exact location in the alignment and refer to the same coordinates on the CDD domains or PFAM HMM profiles; protein domains were not allowed to overlap, and all protein domains combined of a blast hit had to be longer than 300 bp. Note that protein annotation by HMMER and RPSTBLASTN only include gaps of multitudes of three nucleotides, which eliminates the possibility of a frameshift mutation being included in a predicted protein annotation. In the case of multiple TE-related protein domains passing these filters for a single candidate HTT, the alignments were concatenated to obtain a single Ks value per candidate HTT. If the estimated *Ks* values of the candidate HTT were smaller than the 0.5% quantile of the *Ks* distribution of BUSCO genes of the most recent common ancestor of the two involved fungal genomes, the TE pair was kept for further analyses and is hereafter referred to as candidate HTT. This approach yielded 1,312,535 candidate HTTs in 2,639 clade pairs, out of 611,065 possible clade pairs (1,106*1,105/2). Out of 1,905,761 TE copies suitable for HTT detection, 3.4% is involved in HTT (64,814 TE copies).

### Tc1/Mariner alignment in Calycina marina and Lipomyces starkeyi

An alignment 10Kb up and downstream of a candidate HTT involving two DNA *Tc1/Mariner* transposons located in *C. marina* and *L. starkeyi* genomes, belonging to Leotiomycetes and Saccharomycotina, respectively, revealed only an alignment involving the TEs (NUCmer^157^, version 3.1), visualized using the GGGenomes^158^ (1.0.1) package in R.

### Clustering of HTT candidates

Since not all candidate HTTs represent independent transfers, we applied a clustering approach to determine independent transfers per fungal clade pair. First, to obtain percent identity values for TEs within a clade, an all-vs-all BLASTn (-dbsize 10,000,000,000 -evalue 10 -max_target_seqs 600,000) search was performed. Importantly, we only kept hits within a single clade. Next, for each pair of clades, all candidate HTTs between both clades were treated as nodes in a graph. Candidate HTTs were connected if they involved the same TE copy or if the between species percentage identity of involved TEs was greater than the within species percent identity of involved TE (obtained with the BLASTn search, as described above). Next, the cluster_fast_greedy algorithm^159^ from the igraph package (v0.11.8)^160^ was applied in Python (v3.12.3) to detect hit communities. Hit communities having more than 5% of possible edges between them realized were merged (communities with a connectivity greater than 0.05), to prioritize minimizing false positives over false negatives. For computational efficiency, community detection for three clade pairs sharing more than 80,000 candidate HTTs and 3 billion edges was performed using the Leiden algorithm implemented in C++ in the leidenalg package (v0.10.2)^161^, which relies on the igraph package (v0.11.8)^160^ in Python (v3.12.8). This approach yielded 11,001 hit communities, with a minimum of two and a maximum of 3,908 TEs in a hit community.

### Counting minimal number of HTT events

Theoretically, horizontal transfers could have involved a recent common ancestor and a distantly related species. In the hit communities detected above, those would then be counted double, as descendants of this recent common ancestor each have an independent transfer with the distantly related species. To take this into account, we created a network with communities as nodes and edges between communities that share TEs; note that this network does include all clade pairs. To be highly stringent, single linkage clustering was applied to detect clusters using the networkX package (v2.5.1)^162^ in Python (v3.6.13). For each cluster, the species tree was trimmed to only contain the species and collapsed nodes present in the cluster using the ete3 package (v3.1.13)^145^, implemented in Python (v3.6.13). Collapsed nodes were treated as a single taxonomic entity. Per node in the tree, if multiple transfers occurred involving different clades, the minimal number of horizontal transfers needed to explain the observed pattern parsimoniously is 1. In case of multiple independent transfers involving the two same clades (independent transfers have been established per clade pair in the “*Clustering of HTT candidates*” section above), that exact number of independent transfers would be counted for that node. In the end, all counts per node were summed up, which led to a conservative estimate of a minimum number of 5,518 HTT events.

### Enrichment analysis of taxonomic groups and TE types

To determine enrichment of taxonomic groups or TE types, all TEs possessing TE-related protein domains (1,905,761 TEs) in 1,348 fungal genome assemblies were considered. Each genome assemblies was assigned to specific taxonomic groups (**Table S1**). To test whether the TEs present in that taxonomic group were enriched in HTTs, we conducted a Fisher’s exact test for enrichment in R (v4.4.3), where counts in each contingency table represented the number of TEs involved or not involved in HTT per taxonomic group. We also conducted an analogous enrichment analysis, considering TE superfamilies. We also applied these enrichment analyses on the level of individual genomes. To correct for multiple testing, we applied the Benjamin-Hochberg false discovery rate correction for all tests performed.

### Identification of proportion HTT-associated TEs in mobilomes

To identify the proportion of TEs (including fragments) in mobilomes that are descendants of a horizontally transferred TEs, we clustered TEs per genome assembly, based on sequence similarity using MMseqs2 (15.6f452, easy-cluster -c 0.2 --cov-mode 1 –min-seq-id [percent identity threshold])^163^; for each hit community (independent horizontal transfer event), we used the lowest percent identity of the involved candidate HTTs as minimal sequence identity threshold in MMseqs2. We did not distinguish between HTT acceptors and donors, and consequently HTT-associated TEs include descendants of the TEs transferred to the acceptor’s genome, and descendants of TEs that remained in the donor’s genome.

### TE defense mechanisms

Repeat-Induced Point (RIP) mutation pathway d is considered an important genome defense process. RIP specifically induces CpT to TpT and ApG to ApA mutations. We detected RIP signatures per TE copy (*n* = 16,410,702) by calculating their composite RIP-index^164^, where a positive composite RIP-index indicates RIP activity. We classified genomes in RIP-proficient or deficient based on whether > 40% of repeats showed signals of RIP activity.

We also identified five key enzymes involved in TE defenses using DIAMOND^165^ (v2.1.11.165) in sensitive mode, identifying bidirectional best hits. The protein sequence for DNA (cytosine-5-)-methyltransferase (DNMT5) of *Cryptococcus neoformans* (DNMT5) and DNA (cytosine-5-)-methyltransferase (DIM-2), histone-lysine methyltransferase (DIM-5), putative cytosine methyltransferase (RID; stands for RIp Defective), and set-domain histone methylgransferase-7 (SET7) in *Neurospora crassa* were downloaded from the U.S. Department of Energy (DOE) Joint Genome Institute (JGI)^68^ in April 2025.

### Genome compartmentalization

We assessed genome compartmentalized by the presence/absence of TE-rich and gene-sparse regions. For each genome, we calculated per gene the distances to neighboring genes and to the nearest TE. Genomes were considered compartmentalized if >5% of genes had neighboring genes located > 5 kb away and a TE located within 1 kb.

To assess the genomic context of horizontally transferred TEs, we calculated the distances from each horizontally transferred TE to the nearest gene, orphan gene, and effector gene. Orphan genes were defined as singletons – genes that did not cluster with any other protein-coding genes in the 1,348 fungal genome assemblies – based on sequence similarity clustering performed with MMseqs2 (14.7e284, easy-cluster -c 0.8 --cov-mode 1 --min-seq-id 0.5)^163^. Effector genes were identified in two steps: first, we predicted secreted proteins from the protein-coding genes using SignalP (version 4.0)^166^; second, effector genes were predicted among the secreted protein using EffectorP (v3.0)^167^.

## Acknowledgements

This research was financially supported by the Incentive Grant of Utrecht University. (A proportion of) These data were produced by the US Department of Energy Joint Genome Institute in collaboration with the user community. We would like to acknowledge Jan Kees van Amerongen for his invaluable support, Bas Odekerken for his help with analyzing protein sequences, as well as Berend Snel and Daniel Tamarit for their feedback on the initial draft of this manuscript.

## References

1. Gupta, Y. K. et al. Major proliferation of transposable elements shaped the genome of the soybean rust pathogen *Phakopsora pachyrhizi* . Nat. Commun. 14, 1835 (2023).

2. Meyer, A. et al. Giant lungfish genome elucidates the conquest of land by vertebrates. Nature 590, 284–289 (2021).

3. Orgel, L. E. & Crick, F. H. C. Selfish DNA: the ultimate parasite. Nature 284, 604–607 (1980).

4. Doolittle, W. F. & Sapienza, C. Selfish genes, the phenotype paradigm and genome evolution. Nature 284, 601–603 (1980).

5. Badet, T., Feurtey, A. & Croll, D. Recent reactivation of a pathogenicity-associated transposable element triggers major chromosomal rearrangements in a fungal wheat pathogen. 2023.03.29.534637 Preprint at 10.1101/2023.03.29.534637 (2023).

6. Eichler, E. E. & Sankoff, D. Structural dynamics of eukaryotic chromosome evolution. Science 301, 793–797 (2003).

7. Fueyo, R., Judd, J., Feschotte, C. & Wysocka, J. Roles of transposable elements in the regulation of mammalian transcription. Nat. Rev. Mol. Cell Biol. 23, 481–497 (2022).

8. Villanueva-Cañas, J. L., Horvath, V., Aguilera, L. & González, J. Diverse families of transposable elements affect the transcriptional regulation of stress-response genes in *Drosophila melanogaster*. Nucleic Acids Res. 47, 6842–6857 (2019).

9. Wang, C., Milgate, A. W., Solomon, P. S. & McDonald, M. C. The identification of a transposon affecting the asexual reproduction of the wheat pathogen *Zymoseptoria tritici*. Mol. Plant Pathol. 22, 800–816 (2021).

10. Kofler, R., Senti, K.-A., Nolte, V., Tobler, R. & Schlötterer, C. Molecular dissection of a natural transposable element invasion. Genome Res. 28, 824–835 (2018).

11. Zhou, W., Liang, G., Molloy, P. L. & Jones, P. A. DNA methylation enables transposable element-driven genome expansion. Proc. Natl. Acad. Sci. 117, 19359–19366 (2020).

12. Wells, J. N. & Feschotte, C. A field guide to eukaryotic transposable elements. Annu. Rev. Genet. 54, 539–561 (2020).

13. Zeng, X. et al. Comparative analysis of transposable element evolution in crustaceans. Genome Biol. Evol. evaf115 (2025) doi:10.1093/gbe/evaf115.

14. Wang, S. et al. MBD2 couples DNA methylation to transposable element silencing during male gametogenesis. Nat. Plants 10, 13–24 (2024).

15. Huang, Y., et al. Polymorphic transposable elements contribute to variation in recombination landscapes. Proc. Natl. Acad. Sci. 122, e2427312122 (2025).

16. Jeseničnik, T., Štajner, N., Radišek, S. & Jakše, J. RNA interference core components identified and characterised in *Verticillium nonalfalfae*, a vascular wilt pathogenic plant fungi of hops. Sci. Rep. 9, 8651 (2019).

17. Villalba de la Peña, M., Summanen, P. A. M., Liukkonen, M. & Kronholm, I. Chromatin structure influences rate and spectrum of spontaneous mutations in *Neurospora crassa*. Genome Res. 33, 599– 611 (2023).

18. Makova, K. D. & Hardison, R. C. The effects of chromatin organization on variation in mutation rates in the genome. Nat. Rev. Genet. 16, 213–223 (2015).

19. Liu, P., Cuerda-Gil, D., Shahid, S. & Slotkin, R. K. The epigenetic control of the transposable element life cycle in plant genomes and beyond. Annu. Rev. Genet. 56, 63–87 (2022).

20. Ramakrishna, W. et al. Different types and rates of genome evolution detected by comparative sequence analysis of orthologous segments from four cereal genomes. Genetics 162, 1389–1400 (2002).

21. Robillard, É., Le Rouzic, A., Zhang, Z., Capy, P. & Hua-Van, A. Experimental evolution reveals hyperparasitic interactions among transposable elements. Proc. Natl. Acad. Sci. 113, 14763–14768 (2016).

22. Langmüller, A. M., Nolte, V., Dolezal, M. & Schlötterer, C. The genomic distribution of transposable elements is driven by spatially variable purifying selection. Nucleic Acids Res. 51, 9203–9213 (2023).

23. Schaack, S., Gilbert, C. & Feschotte, C. Promiscuous DNA: horizontal transfer of transposable elements and why it matters for eukaryotic evolution. Trends Ecol. Evol. 25, 537–546 (2010).

24. Bourque, G. et al. Ten things you should know about transposable elements. Genome Biol. 19, 199 (2018).

25. Daniels, S. B., Peterson, K. R., Strausbaugh, L. D., Kidwell, M. G. & Chovnick, A. Evidence for horizontal transmission of the *P* transposable element between *Drosophila* species. Genetics 124, 339– 355 (1990).

26. Venner, S. et al. Ecological networks to unravel the routes to horizontal transposon transfers. PLOS Biol. 15, e2001536 (2017).

27. Peccoud, J., Loiseau, V., Cordaux, R. & Gilbert, C. Massive horizontal transfer of transposable elements in insects. Proc. Natl. Acad. Sci. 114, 4721–4726 (2017).

28. Dunemann, S. M. & Wasmuth, J. D. Horizontal transfer of a retrotransposon between parasitic nematodes and the common shrew. Mob. DNA 10, 24 (2019).

29. Widen, S. A. et al. Virus-like transposons cross the species barrier and drive the evolution of genetic incompatibilities. Science 380, eade0705 (2023).

30. Urquhart, A. S., O’Donnell, S., Gluck-Thaler, E. & Vogan, A. A. A natural mechanism of eukaryotic horizontal gene transfer. 2025.02.28.640899 Preprint at 10.1101/2025.02.28.640899 (2025).

31. Diao, X., Freeling, M. & Lisch, D. Horizontal transfer of a plant transposon. PLOS Biol. 4, e5 (2005).

32. Walsh, A. M., Kortschak, R. D., Gardner, M. G., Bertozzi, T. & Adelson, D. L. Widespread horizontal transfer of retrotransposons. Proc. Natl. Acad. Sci. 110, 1012–1016 (2013).

33. Ivancevic, A. M., Kortschak, R. D., Bertozzi, T. & Adelson, D. L. Horizontal transfer of *BovB* and *L1* retrotransposons in eukaryotes. Genome Biol. 19, 85 (2018).

34. Galbraith, J. D., Ludington, A. J., Sanders, K. L., Suh, A. & Adelson, D. L. Horizontal transfer and subsequent explosive expansion of a DNA transposon in sea kraits (*Laticauda*). Biol. Lett. 17, 20210342 (2021).

35. Zhang, H.-H., Peccoud, J., Xu, M.-R.-X., Zhang, X.-G. & Gilbert, C. Horizontal transfer and evolution of transposable elements in vertebrates. Nat. Commun. 11, 1362 (2020).

36. Muller, H., Savisaar, R., Peccoud, J., Charlat, S. & Gilbert, C. Phylogenetic relatedness rather than aquatic habitat fosters horizontal transfer of transposable elements in animals. 2024.12.18.629015 Preprint at 10.1101/2024.12.18.629015 (2024).

37. Pianezza, R. & Kofler, R. Biogeography shapes the TE landscape of Drosophila melanogaster. 2025.05.22.655554 Preprint at 10.1101/2025.05.22.655554 (2025).

38. Blackwell, M. The fungi: 1, 2, 3 5.1 million species? Am. J. Bot. 98, 426–438 (2011).

39. Naranjo-Ortiz, M. A. & Gabaldón, T. Fungal evolution: major ecological adaptations and evolutionary transitions. Biol. Rev. Camb. Philos. Soc. 94, 1443–1476 (2019).

40. de Mattos-Shipley, K. M. J. et al. The good, the bad and the tasty: The many roles of mushrooms. Stud. Mycol. 85, 125–157 (2016).

41. Zeilinger, S. et al. Friends or foes? Emerging insights from fungal interactions with plants. FEMS Microbiol. Rev. 40, 182–207 (2016).

42. Meyer, V. et al. Current challenges of research on filamentous fungi in relation to human welfare and a sustainable bio-economy: a white paper. Fungal Biol. Biotechnol. 3, 6 (2016).

43. Lorrain, C., Feurtey, A., Möller, M., Haueisen, J. & Stukenbrock, E. Dynamics of transposable elements in recently diverged fungal pathogens: lineage-specific transposable element content and efficiency of genome defenses. G3 GenesGenomesGenetics 11, jkab068 (2021).

44. Grandaubert, J., Balesdent, M.-H. & Rouxel, T. Chapter Three - Evolutionary and Adaptive Role of Transposable Elements in Fungal Genomes. in Advances in Botanical Research (ed. Martin, F. M.) vol. 70 79–107 (Academic Press, 2014).

45. Castanera, R. et al. Transposable elements versus the fungal genome: impact on whole-genome architecture and transcriptional profiles. PLoS Genet. 12, e1006108 (2016).

46. Díaz, C. L. et al. Transposons and accessory genes drive adaptation in a clonally evolving fungal pathogen. 2025.02.18.635021 Preprint at 10.1101/2025.02.18.635021 (2025).

47. Oggenfuss, U., Badet, T. & Croll, D. A systematic screen for co-option of transposable elements across the fungal kingdom. Mob. DNA 15, 2 (2024).

48. Glass, N. L. & Kaneko, I. Fatal attraction: nonself recognition and heterokaryon incompatibility in filamentous fungi. Eukaryot. Cell 2, 1–8 (2003).

49. Jakobsen, I. Hyphal fusion to plant species connections – giant mycelia and community nutrient flow. New Phytol. 164, 4–7 (2004).

50. Shoji, J., Charlton, N. D., Yi, M., Young, C. A. & Craven, K. D. Vegetative hyphal fusion and subsequent nuclear behavior in epichloë grass endophytes. PLoS ONE 10, e0121875 (2015).

51. Novais, C. B. de, Pepe, A., Siqueira, J. O., Giovannetti, M. & Sbrana, C. Compatibility and incompatibility in hyphal anastomosis of arbuscular mycorrhizal fungi. Sci. Agric. 74, 411–416 (2017).

52. Craven, K. D., Vélëz, H., Cho, Y., Lawrence, C. B. & Mitchell, T. K. Anastomosis is required for virulence of the fungal necrotroph *Alternaria brassicicola*. Eukaryot. Cell 7, 675–683 (2008).

53. O’Connor, E. et al. Proteomic investigation of interhyphal interactions between strains of *Agaricus bisporus*. Fungal Biol. 124, 579–591 (2020).

54. Giovannetti, M., Azzolini, D. & Citernesi, A. S. Anastomosis formation and nuclear and protoplasmic exchange in arbuscular mycorrhizal fungi. Appl. Environ. Microbiol. 65, 5571–5575 (1999).

55. Hill, R. et al. Starship giant transposable elements cluster by host taxonomy using k-mer-based phylogenetics. G3 GenesGenomesGenetics jkaf082 (2025) doi:10.1093/g3journal/jkaf082.

56. Bucknell, A. et al. Sanctuary: A Starship transposon facilitating the movement of the virulence factor ToxA in fungal wheat pathogens. 2024.03.04.583430 Preprint at 10.1101/2024.03.04.583430 (2024).

57. Urquhart, A., Vogan, A. A. & Gluck-Thaler, E. *Starships*: a new frontier for fungal biology. Trends Genet. 40, 1060–1073 (2024).

58. Wallau, G. L., Ortiz, M. F. & Loreto, E. L. S. Horizontal transposon transfer in Eukarya: detection, bias, and perspectives. Genome Biol. Evol. 4, 801–811 (2012).

59. Stajich, J. E. Fungal genomes and insights into the evolution of the kingdom. Microbiol. Spectr. 5, (2017).

60. Harder, C. B. et al. Extreme overall mushroom genome expansion in *Mycena* s.s. irrespective of plant hosts or substrate specializations. Cell Genomics 4, 100586 (2024).

61. Haridas, S. et al. 101 Dothideomycetes genomes: A test case for predicting lifestyles and emergence of pathogens. Stud. Mycol. 96, 141–153 (2020).

62. Zaccaron, A. Z. & Stergiopoulos, I. The dynamics of fungal genome organization and its impact on host adaptation and antifungal resistance. J. Genet. Genomics (2024) doi:10.1016/j.jgg.2024.10.010.

63. Corre, E., Morin, E., Duplessis, S. & Lorrain, C. Ancestral and recent bursts of transposition shaped the massive genomes of plant pathogenic rust fungi. 2025.01.10.632365 Preprint at 10.1101/2025.01.10.632365 (2025).

64. Tobias, P. A., et al. *Austropuccinia psidii*, causing myrtle rust, has a gigabase-sized genome shaped by transposable elements. G3 Bethesda Md 11, jkaa015 (2021).

65. Stajich, J. E. et al. Signatures of transposon-mediated genome inflation, host specialization, and photoentrainment in *Entomophthora muscae* and allied entomophthoralean fungi. eLife 12, RP92863 (2024).

66. Grandaubert, J. et al. Transposable element-assisted evolution and adaptation to host plant within the *Leptosphaeria maculans-Leptosphaeria biglobosa* species complex of fungal pathogens. BMC Genomics 15, 891 (2014).

67. Ohm, R. A. et al. Diverse lifestyles and strategies of plant pathogenesis encoded in the genomes of eighteen Dothideomycetes fungi. PLOS Pathog. 8, e1003037 (2012).

68. Nordberg, H. et al. The genome portal of the Department of Energy Joint Genome Institute: 2014 updates. Nucleic Acids Res. 42, D26–31 (2014).

69. Tedersoo, L. et al. High-level classification of the Fungi and a tool for evolutionary ecological analyses. Fungal Divers. 90, 135–159 (2018).

70. Wicker, T. et al. A unified classification system for eukaryotic transposable elements. Nat. Rev. Genet. 8, 973–982 (2007).

71. Selker, E. U. & Garrett, P. W. DNA sequence duplications trigger gene inactivation in *Neurospora crassa*. Proc. Natl. Acad. Sci. 85, 6870–6874 (1988).

72. Selker, E. U. et al. The methylated component of the *Neurospora crassa* genome. Nature 422, 893– 897 (2003).

73. van Wyk, S., Wingfield, B. D., De Vos, L., van der Merwe, N. A. & Steenkamp, E. T. Genome-wide analyses of repeat-induced point mutations in the Ascomycota. Front. Microbiol. 11, (2021).

74. He, Z. et al. RID is required for both repeat-induced point mutation and nucleation of a novel transitional heterochromatic state for euchromatic repeats. Nucleic Acids Res. 53, gkaf263 (2025).

75. Lewis, Z. A. et al. DNA methylation and normal chromosome behavior in *Neurospora* depend on five components of a histone methyltransferase complex, DCDC. PLoS Genet. 6, e1001196 (2010).

76. Jamieson, K., Rountree, M. R., Lewis, Z. A., Stajich, J. E. & Selker, E. U. Regional control of histone H3 lysine 27 methylation in *Neurospora*. Proc. Natl. Acad. Sci. 110, 6027–6032 (2013).

77. Jeltsch, A., Nellen, W. & Lyko, F. Two substrates are better than one: dual specificities for Dnmt2 methyltransferases. Trends Biochem. Sci. 31, 306–308 (2006).

78. Catania, S. et al. Evolutionary persistence of DNA methylation for millions of years after ancient loss of a de novo methyltransferase. Cell 180, 263–277.e20 (2020).

79. Kouzminova, E. & Selker, E. U. dim-2 encodes a DNA methyltransferase responsible for all known cytosine methylation in *Neurospora*. EMBO J. 20, 4309–4323 (2001).

80. John Clutterbuck, A. Genomic evidence of repeat-induced point mutation (RIP) in filamentous ascomycetes. Fungal Genet. Biol. 48, 306–326 (2011).

81. Horns, F., Petit, E., Yockteng, R. & Hood, M. E. Patterns of repeat-induced point mutation in transposable elements of basidiomycete fungi. Genome Biol. Evol. 4, 240–247 (2012).

82. Badet, T. & Croll, D. A systematic screen for genetic factors underpinning transposon defense systems across the fungal kingdom. 2025.01.10.632494 Preprint at 10.1101/2025.01.10.632494 (2025).

83. Lorrain, C., Feurtey, A., Möller, M., Haueisen, J. & Stukenbrock, E. Dynamics of transposable elements in recently diverged fungal pathogens: lineage-specific transposable element content and efficiency of genome defenses. G3 GenesGenomesGenetics 11, jkab068 (2021).

84. Cuomo, C. A. et al. The *Fusarium graminearum* genome reveals a link between localized polymorphism and pathogen specialization. Science 317, 1400–1402 (2007).

85. Graïa, F. et al. Genome quality control: RIP (repeat-induced point mutation) comes to Podospora. Mol. Microbiol. 40, 586–595 (2001).

86. Ikeda, K. et al. Repeat-induced point mutation (RIP) in *Magnaporthe grisea*: implications for its sexual cycle in the natural field context. Mol. Microbiol. 45, 1355–1364 (2002).

87. Idnurm, A. & Howlett, B. Analysis of loss of pathogenicity mutants reveals that repeat-induced point mutations can occur in the Dothideomycete Leptosphaeria maculans. Fungal Genet. Biol. FG B 39, (2003).

88. Coleman, J. J. et al. The genome of *Nectria haematococca*: contribution of supernumerary chromosomes to gene expansion. PLOS Genet. 5, e1000618 (2009).

89. Pomraning, K. R., Connolly, L. R., Whalen, J. P., Smith, K. M. & Freitag, M. Repeat-induced point mutation, DNA methylation and heterochromatin in Gibberella zeae (Anamorph: Fusarium graminearum). in Fusarium Genomics, Molecular and Cellular Biology 93–109 (Caister Academic Press, Great Britain).

90. Dong, S., Raffaele, S. & Kamoun, S. The two-speed genomes of filamentous pathogens: waltz with plants. Curr. Opin. Genet. Dev. 35, 57–65 (2015).

91. Wacker, T., et al. Two-speed genome evolution drives pathogenicity in fungal pathogens of animals. Proc. Natl. Acad. Sci. 120, e2212633120 (2023).

92. Rouxel, T. et al. Effector diversification within compartments of the *Leptosphaeria maculans* genome affected by Repeat-Induced Point mutations. Nat. Commun. 2, 202 (2011).

93. Kim, K.-T. et al. Evolution of the genes encoding effector candidates within multiple pathotypes of *Magnaporthe oryzae*. Front. Microbiol. 10, 2575 (2019).

94. van Dam, P. et al. A mobile pathogenicity chromosome in *Fusarium oxysporum* for infection of multiple cucurbit species. Sci. Rep. 7, 9042 (2017).

95. Duplessis, S. et al. Obligate biotrophy features unraveled by the genomic analysis of rust fungi. Proc. Natl. Acad. Sci. U. S. A. 108, 9166–9171 (2011).

96. Feurtey, A. et al. Genome compartmentalization predates species divergence in the plant pathogen genus Zymoseptoria. BMC Genomics 21, 588 (2020).

97. Cook, D. E., Kramer, H. M., Torres, D. E., Seidl, M. F. & Thomma, B. P. H. J. A unique chromatin profile defines adaptive genomic regions in a fungal plant pathogen. eLife 9, e62208 (2020).

98. Engelbrecht, J., Duong, T. A., Prabhu, S. A., Seedat, M. & van den Berg, N. Genome of the destructive oomycete *Phytophthora cinnamomi* provides insights into its pathogenicity and adaptive potential. BMC Genomics 22, 302 (2021).

99. Torres, D. E., Thomma, B. P. H. J. & Seidl, M. F. Transposable elements contribute to genome dynamics and gene expression variation in the fungal plant pathogen *Verticillium dahliae*. Genome Biol. Evol. 13, evab135 (2021).

100. Torres, D. E., Oggenfuss, U., Croll, D. & Seidl, M. F. Genome evolution in fungal plant pathogens: looking beyond the two-speed genome model. Fungal Biol. Rev. 34, 136–143 (2020).

101. Wang, Q., Wang, Y., Wang, J., Gong, Z. & Han, G.-Z. Plants acquired a major retrotransposon horizontally from fungi during the conquest of land. New Phytol. 232, 11–16 (2021).

102. Bergman, C. M. Horizontal transfer and proliferation of *Tsu4* in *Saccharomyces paradoxus*. Mob. DNA 9, 18 (2018).

103. Sato, Y. et al. Starship giant transposons dominate plastic genomic regions in a fungal plant pathogen and drive virulence evolution. 2025.01.08.631984 Preprint at 10.1101/2025.01.08.631984 (2025).

104. Urquhart, A. S., Chong, N. F., Yang, Y. & Idnurm, A. A large transposable element mediates metal resistance in the fungus *Paecilomyces variotii*. Curr. Biol. CB 32, 937–950.e5 (2022).

105. Gluck-Thaler, E. & Vogan, A. A. Systematic identification of cargo-mobilizing genetic elements reveals new dimensions of eukaryotic diversity. Nucleic Acids Res. 52, 5496–5513 (2024).

106. Shi, S. et al. Prokaryotic and eukaryotic horizontal transfer of sailor (*DD82E*), a new superfamily of *IS630-Tc1-mariner* DNA transposons. Biology 10, 1005 (2021).

107. Orozco-Arias, S., Dupeyron, M., Gutiérrez-Duque, D., Tabares-Soto, R. & Guyot, R. High nucleotide similarity of three *Copia* lineage LTR retrotransposons among plant genomes. Genome 66, 51–61 (2023).

108. Park, M., Sarkhosh, A., Tsolova, V. & El-Sharkawy, I. Horizontal transfer of LTR retrotransposons contributes to the genome diversity of Vitis. Int. J. Mol. Sci. 22, 10446 (2021).

109. El Baidouri, M., et al. Widespread and frequent horizontal transfers of transposable elements in plants. Genome Res. 24, 831–838 (2014).

110. Bartolomé, C., Bello, X. & Maside, X. Widespread evidence for horizontal transfer of transposable elements across Drosophila genomes. Genome Biol. 10, R22 (2009).

111. Lange, B., Kaufmann, A. P. & Ebert, D. Genetic, ecological and geographic covariables explaining host range and specificity of a microsporidian parasite. J. Anim. Ecol. 84, 1711–1719 (2015).

112. Longdon, B., Brockhurst, M. A., Russell, C. A., Welch, J. J. & Jiggins, F. M. The evolution and genetics of virus host shifts. PLOS Pathog. 10, e1004395 (2014).

113. Perlman, S. J. & Jaenike, J. Infection success in novel hosts: an experimental and phylogenetic study of *Drosophila*-parasitic nematodes. Evolution 57, 544–557 (2003).

114. Wells, K. & Clark, N. J. Host specificity in variable environments. Trends Parasitol. 35, 452–465 (2019).

115. Gilbert, C. & Feschotte, C. Horizontal acquisition of transposable elements and viral sequences: patterns and consequences. Curr. Opin. Genet. Dev. 49, 15–24 (2018).

116. Peccoud, J., Cordaux, R. & Gilbert, C. Analyzing horizontal transfer of transposable elements on a large scale: challenges and prospects. BioEssays 40, 1700177 (2018).

117. Morogovsky, A., Handelman, M., Abou Kandil, A., Shadkchan, Y. & Osherov, N. Horizontal gene transfer of triazole resistance in *Aspergillus fumigatus*. Microbiol. Spectr. 10, e01112–22 (2022).

118. Ruiz-Roldán, M. C. et al. Nuclear dynamics during germination, conidiation, and hyphal fusion of *Fusarium oxysporum*. Eukaryot. Cell 9, 1216–1224 (2010).

119. Silva, J. C., Loreto, E. L. & Clark, J. B. Factors that affect the horizontal transfer of transposable elements. Curr. Issues Mol. Biol. 6, 57–71 (2004).

120. Habig, M., et al. Frequent horizontal chromosome transfer between asexual fungal insect pathogens. Proc. Natl. Acad. Sci. 121, e2316284121 (2024).

121. Deng, Y. et al. Viral cross-class transmission results in disease of a phytopathogenic fungus. ISME J. 16, 2763–2774 (2022).

122. Fischer, M. G. & Suttle, C. A. A virophage at the origin of large DNA transposons. Science 332, 231–234 (2011).

123. Krupovic, M., Bamford, D. H. & Koonin, E. V. Conservation of major and minor jelly-roll capsid proteins in Polinton (Maverick) transposons suggests that they are bona fide viruses. Biol. Direct 9, 6 (2014).

124. Pastuzyn, E. D. et al. The neuronal gene arc encodes a repurposed retrotransposon Gag protein that mediates intercellular RNA transfer. Cell 172, 275–288.e18 (2018).

125. Kim, A. et al. Retroviruses in invertebrates: the gypsy retrotransposon is apparently an infectious retrovirus of Drosophila melanogaster. Proc. Natl. Acad. Sci. 91, 1285–1289 (1994).

126. Hackl, T. et al. Novel integrative elements and genomic plasticity in ocean ecosystems. Cell 186, 47–62.e16 (2023).

127. Biller, S. J. et al. Distinct horizontal gene transfer potential of extracellular vesicles versus viral-like particles in marine habitats. Nat. Commun. 16, 2126 (2025).

128. Costa, J. H. et al. Phytotoxic tryptoquialanines produced in vivo by *Penicillium digitatum* are exported in extracellular vesicles. mBio 12, 10.1128/mbio.03393-20 (2021).

129. Kwon, S. et al. mRNA Inventory of extracellular vesicles from *Ustilago maydis*. J. Fungi Basel Switz. 7, 562 (2021).

130. Visser, C. et al. Tracking the uptake of labelled host-derived extracellular vesicles by the human fungal pathogen *Aspergillus fumigatus*. microLife 5, uqae022 (2024).

131. Wang, S. et al. Plant mRNAs move into a fungal pathogen via extracellular vesicles to reduce infection. Cell Host Microbe 32, 93–105.e6 (2024).

132. Roth, R. et al. Arbuscular cell invasion coincides with extracellular vesicles and membrane tubules. Nat. Plants 5, 204–211 (2019).

133. Sayers, E. W. et al. GenBank. Nucleic Acids Res. 47, D94–D99 (2019).

134. Schoch, C. L. et al. NCBI Taxonomy: a comprehensive update on curation, resources and tools. Database J. Biol. Databases Curation 2020, baaa062 (2020).

135. Hane, J. K., Paxman, J., Jones, D. A. B., Oliver, R. P. & de Wit, P. “CATAStrophy,” a genome-informed trophic classification of filamentous plant pathogens – how many different types of filamentous plant pathogens are there? Front. Microbiol. 10, (2020).

136. Manni, M., Berkeley, M. R., Seppey, M., Simão, F. A. & Zdobnov, E. M. BUSCO Update: novel and streamlined workflows along with broader and deeper phylogenetic coverage for scoring of eukaryotic, prokaryotic, and viral genomes. Mol. Biol. Evol. 38, 4647–4654 (2021).

137. Manni, M., Berkeley, M. R., Seppey, M. & Zdobnov, E. M. BUSCO: assessing genomic data quality and beyond. Curr. Protoc. 1, e323 (2021).

138. Nguyen, L.-T., Schmidt, H. A., von Haeseler, A. & Minh, B. Q. Iq-tree: a fast and effective stochastic algorithm for estimating maximum-likelihood phylogenies. Mol. Biol. Evol. 32, 268–274 (2015).

139. Deorowicz, S., Debudaj-Grabysz, A. & Gudyś, A. FAMSA: Fast and accurate multiple sequence alignment of huge protein families. Sci. Rep. 6, 33964 (2016).

140. Capella-Gutiérrez, S., Silla-Martínez, J. M. & Gabaldón, T. trimAl: a tool for automated alignment trimming in large-scale phylogenetic analyses. Bioinformatics 25, 1972–1973 (2009).

141. Kalyaanamoorthy, S., Minh, B. Q., Wong, T. K. F., von Haeseler, A. & Jermiin, L. S. ModelFinder: fast model selection for accurate phylogenetic estimates. Nat. Methods 14, 587–589 (2017).

142. Yu, G., Smith, D. K., Zhu, H., Guan, Y. & Lam, T. T.-Y. ggtree: an R package for visualization and annotation of phylogenetic trees with their covariates and other associated data. Methods Ecol. Evol. 8, 28–36 (2017).

143. Parks, D. H. et al. A standardized bacterial taxonomy based on genome phylogeny substantially revises the tree of life. Nat. Biotechnol. 36, 996–1004 (2018).

144. Mölder, F., et al. Sustainable data analysis with Snakemake. Preprint at 10.12688/f1000research.29032.1 (2021).

145. Huerta-Cepas, J., Serra, F. & Bork, P. ETE 3: reconstruction, analysis, and visualization of phylogenomic data. Mol. Biol. Evol. 33, 1635–1638 (2016).

146. Katoh, K. & Standley, D. M. MAFFT multiple sequence alignment software version 7: improvements in performance and usability. Mol. Biol. Evol. 30, 772–780 (2013).

147. Suyama, M., Torrents, D. & Bork, P. PAL2NAL: robust conversion of protein sequence alignments into the corresponding codon alignments. Nucleic Acids Res. 34, W609–W612 (2006).

148. Baril, T., Galbraith, J. & Hayward, A. Earl Grey: a fully automated user-friendly transposable element annotation and analysis pipeline. Mol. Biol. Evol. 41, msae068 (2024).

149. Smit, A., Hubley, R. & Green, P. RepeatMasker Open-4.0. (2013).

150. Hoede, C. et al. PASTEC: an automatic transposable element classification tool. PLOS ONE 9, e91929 (2014).

151. Quinlan, A. R. & Hall, I. M. BEDTools: a flexible suite of utilities for comparing genomic features. Bioinformatics 26, 841–842 (2010).

152. Gryganskyi, A. P. et al. Phylogenetic and phylogenomic definition of *Rhizopus* species. G3 Bethesda Md **8**, 2007–2018 (2018).

153. Muszewska, A., Steczkiewicz, K., Stepniewska-Dziubinska, M. & Ginalski, K. Transposable elements contribute to fungal genes and impact fungal lifestyle. Sci. Rep. 9, 4307 (2019).

154. Rice, P., Longden, I. & Bleasby, A. EMBOSS: the european molecular biology open software suite. Trends Genet. TIG 16, 276–277 (2000).

155. Camacho, C. et al. BLAST+: architecture and applications. BMC Bioinformatics 10, 421 (2009).

156. Marchler-Bauer, A. et al. CDD: NCBI’s conserved domain database. Nucleic Acids Res. 43, D222– D226 (2015).

157. Kurtz, S. et al. Versatile and open software for comparing large genomes. Genome Biol. 5, R12 (2004).

158. Hackl, T., Ankenbrand, M., Adrichem, B. van, Wilkins, D. & Haslinger, K. gggenomes: effective and versatile visualizations for comparative genomics. Preprint at 10.48550/arXiv.2411.13556 (2024).

159. Clauset, A., Newman, M. E. J. & Moore, C. Finding community structure in very large networks. *Phys*. Rev. E 70, 066111 (2004).

160. Csardi, G. & Nepusz, T. The igraph software package for complex network research. Available Igraphorg **Complex Systems**, 1695 (2005).

161. Traag, V. A., Waltman, L. & van Eck, N. J. From Louvain to Leiden: guaranteeing well-connected communities. Sci. Rep. 9, 5233 (2019).

162. Hagberg, A. A., Schult, D. A. & Swart, P. J. Exploring network structure, dynamics, and function using NetworkX. in Proceedings of the 7th Python in Science Conference (eds. Varoquaux, G., Vaught, T. & Millman, J.) 11–15 (Pasadena, CA USA, 2008).

163. Steinegger, M. & Söding, J. MMseqs2 enables sensitive protein sequence searching for the analysis of massive data sets. Nat. Biotechnol. 35, 1026–1028 (2017).

164. van Wyk, S. et al. The RIPper, a web-based tool for genome-wide quantification of Repeat-Induced Point (RIP) mutations. PeerJ 7, e7447 (2019).

165. Buchfink, B., Reuter, K. & Drost, H.-G. Sensitive protein alignments at tree-of-life scale using DIAMOND. Nat. Methods 18, 366–368 (2021).

166. Nielsen, H., Engelbrecht, J., Brunak, S. & von Heijne, G. Identification of prokaryotic and eukaryotic signal peptides and prediction of their cleavage sites. Protein Eng. 10, 1–6 (1997).

167. Sperschneider, J. & Dodds, P. N. EffectorP 3.0: prediction of apoplastic and cytoplasmic effectors in fungi and oomycetes. Mol. Plant-Microbe Interactions® 35, 146–156 (2022).

