## Supplementary Figures for "Extensive horizontal transfer of transposable elements shape fungal mobilomes"

**Supplemental information**

Table S1: Assembly statistics, taxonomy, references, transposable element coverage, and BUSCO scores of genome assemblies used in this study

Table S2: Lifestyle prediction of genome assemblies

Table S3: Transposable element classifications used in this study

Table S4: Transposable element counts per genome

Table S5: Transposable element (TEs) counts per genome (including TEs >300bp and encoding TE-related protein domain)

Table S6: Transposable element (TEs) counts per genome (TEs involved in HTT)

Table S7: Enrichment analyses per transposable element type

Table S8: Enrichments analyses of transposable element types per genome assembly

Table S9: Ka/Ks per transposable element type

Table S10: Ka/Ks per transposable element type, separated by comparison vertically vs horizontally transferred transposable elements

Table S11: Enrichment analyses of fungal taxonomic groups

Table S12: Distances of HTT-derived transposable elements to orphan, effector and regular genes

Table S13: List of recently diverged genomes for which horizontal transfer of transposable elements was not assessed

Table S14: List of -related protein domains

Figure S1: Horizontal transfer of transposable element detection pipeline

Figure S2: Number of genomes enriched or involved in horizontal transfer of transposable elements per transposable element type

Figure S3: Number of hit communities per taxonomic group

Figure S4: Heatmap of number of horizontal transfers of transposable between different lifestyles

Figure S5: Presence and absence of five key enzymes involved in genome defenses

Figure S6: Enrichment analyses for different genome defenses and genome organization

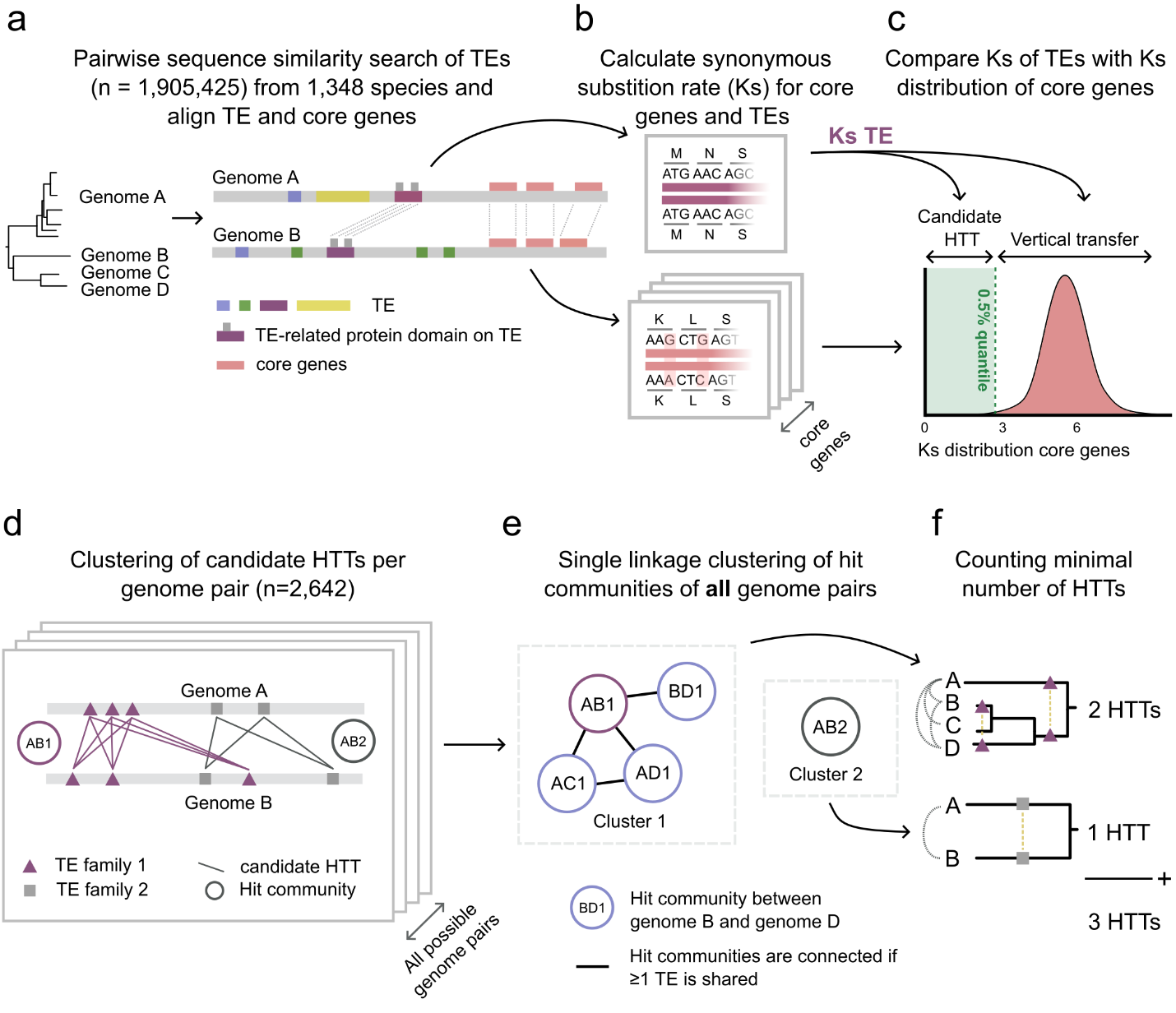

**Figure S1 - Computational approach to systematically detect horizontally transferred transposable elements (HTTs) throughout the fungal kingdom yielded 11,006 independent transfer events. a**, Pairwise sequence similarity searches of transposable elements (TEs) (>300bp and encoding TE-related protein domain, n = 1,905,425) in across 1,348 fungal genomes were performed. TE pairs must cover at least 60% of both TEs. **b**, Synonymous mutation rates were calculated for TE-related protein domains in each TE pair, alongside Ks values for core genes between the corresponding genomes to represent their evolutionary divergence. **c**, we considered a TE pair as a candidate HTT if its Ks value was significantly lower (<0.5% quantile) than expected given the distribution of core genes Ks values. **d**, To identify independent HTT events, TE pairs likely resulting from the same transfer were clustered into hit communities per clade comparison, using the cluster_fast_greedy algorithm^1^ from the igraph package^2^ in python. For graphs with more than 80,000 nodes and 3 million edges, the Leiden algorithm implemented in C++ in the leidenalg package^3^, which relies on the igraph package^2^ in Python was used. TE pairs were connected if they 1) involved the same TE or 2) if their within-species percent identity was higher than any of the between-species percent identity. Subsequently, communities were merged if >5% of edges were realized between them. **e**, Hit communities in all clades (all genomes) were then connected by shared TEs and further grouped using single-linkage clustering to identify clusters of related transfer events. **f**, To avoid overcounting ancestral transfers, each cluster of hit communities was mapped onto the species tree, and we applied a parsimony-based approach to infer the minimal number of independent HTT events required to explain the observed patterns.

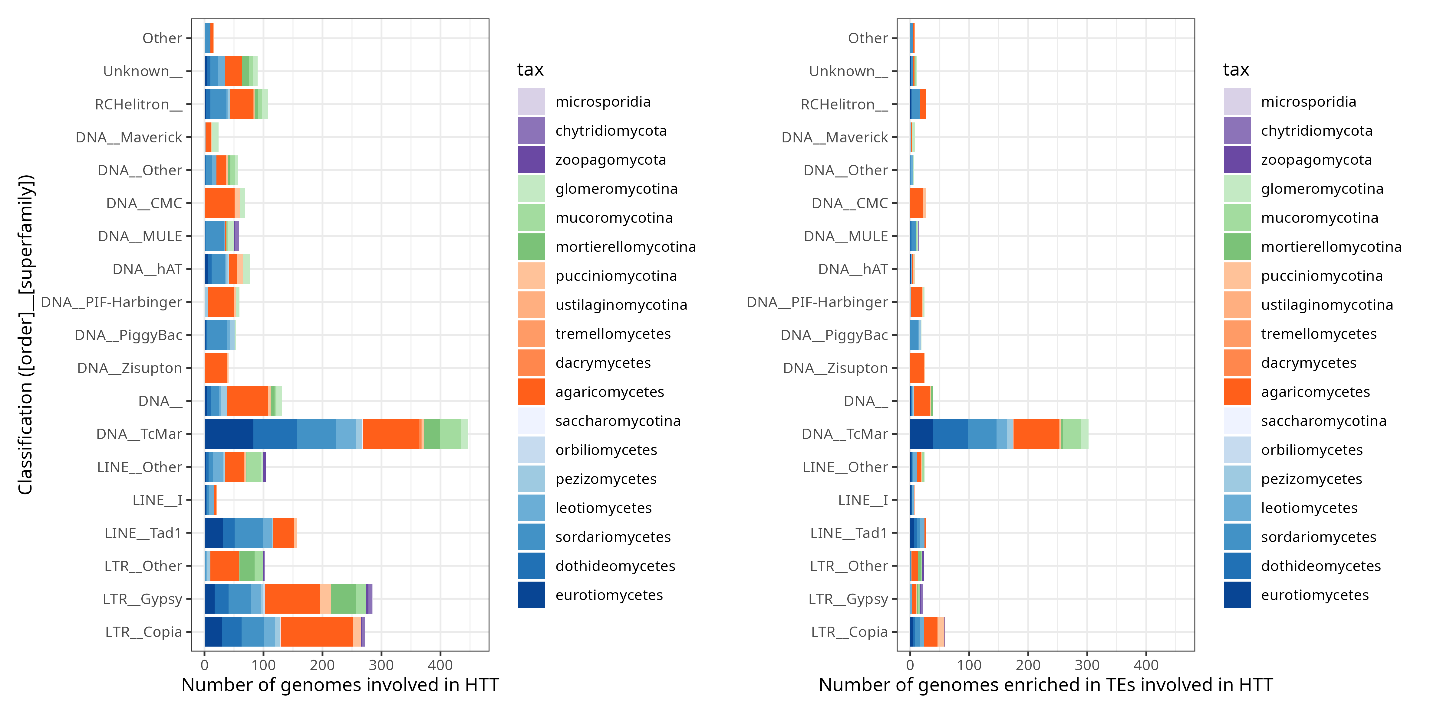

**Figure S2 – Involvement of genomes in horizontal transfer of transposable elements (HTT) and enrichment of transposable elements (TEs) involved in HTT per genome, plotted per TE superfamily and different fungal lineages.** Number of genomes involved in HTT per TE superfamily (left) and number of genomes enriched in TEs involved in HTT per TE superfamily (right), colored by fungal taxonomic lineages. Enrichment tests were performed per genome and TE superfamily using a one-sided Fisher’s exact test, with Bonferroni-Hochberg multiple testing correction (see **Tab. S8**).

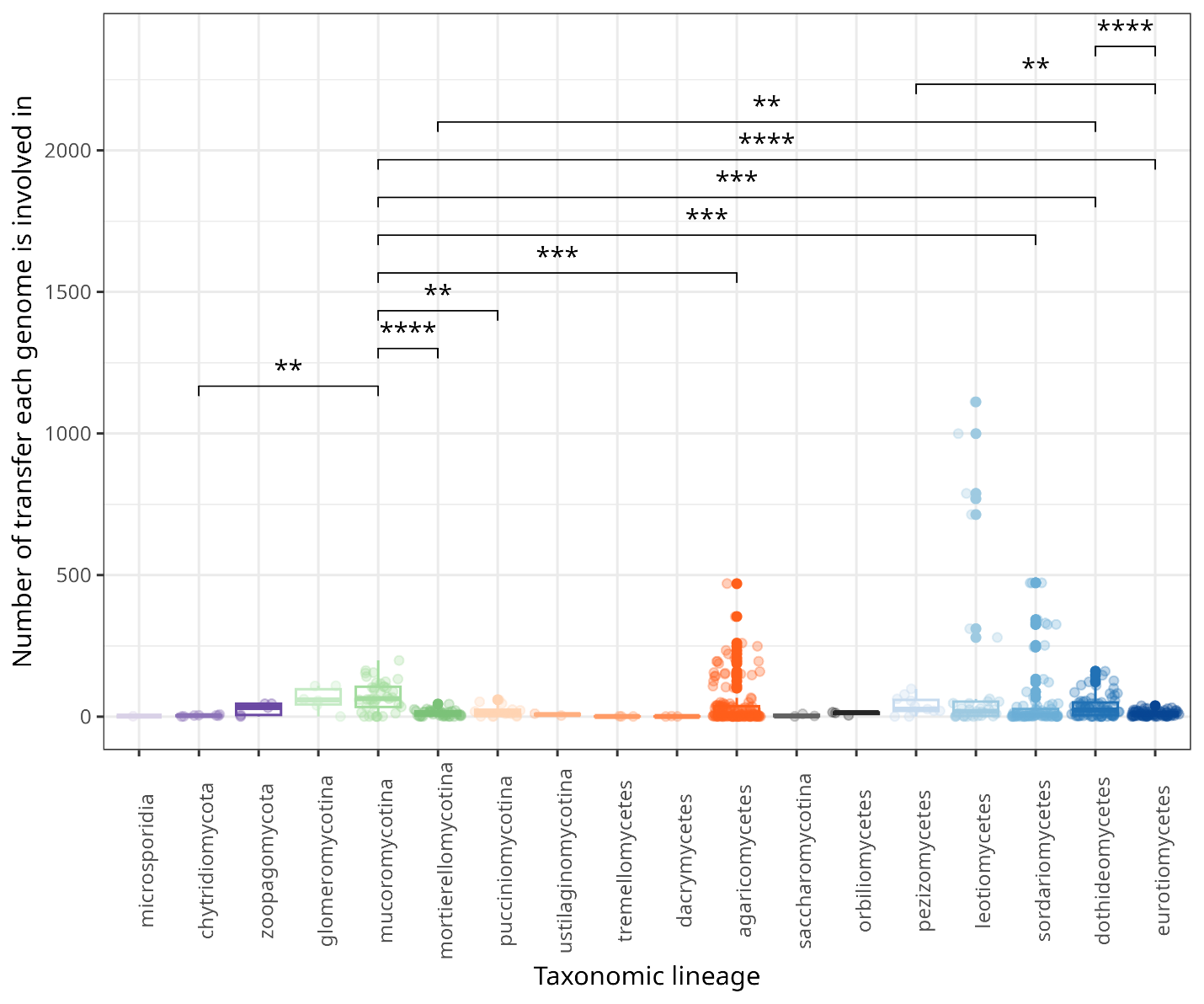

**Figure S3 – The number of horizontal transfers of transposable elements (HTTs) that each genome participates in.** Boxplot of number of HTTs that each genome participates in, measured at the level of hit communities, per fungal lineage. To test a difference in rank totals between fungal lineages, a Kruskal-Wallis test was performed (p = 3.2e-7), and results of subsequent pairwise Wilcoxon rank sum tests are indicated in the plot.

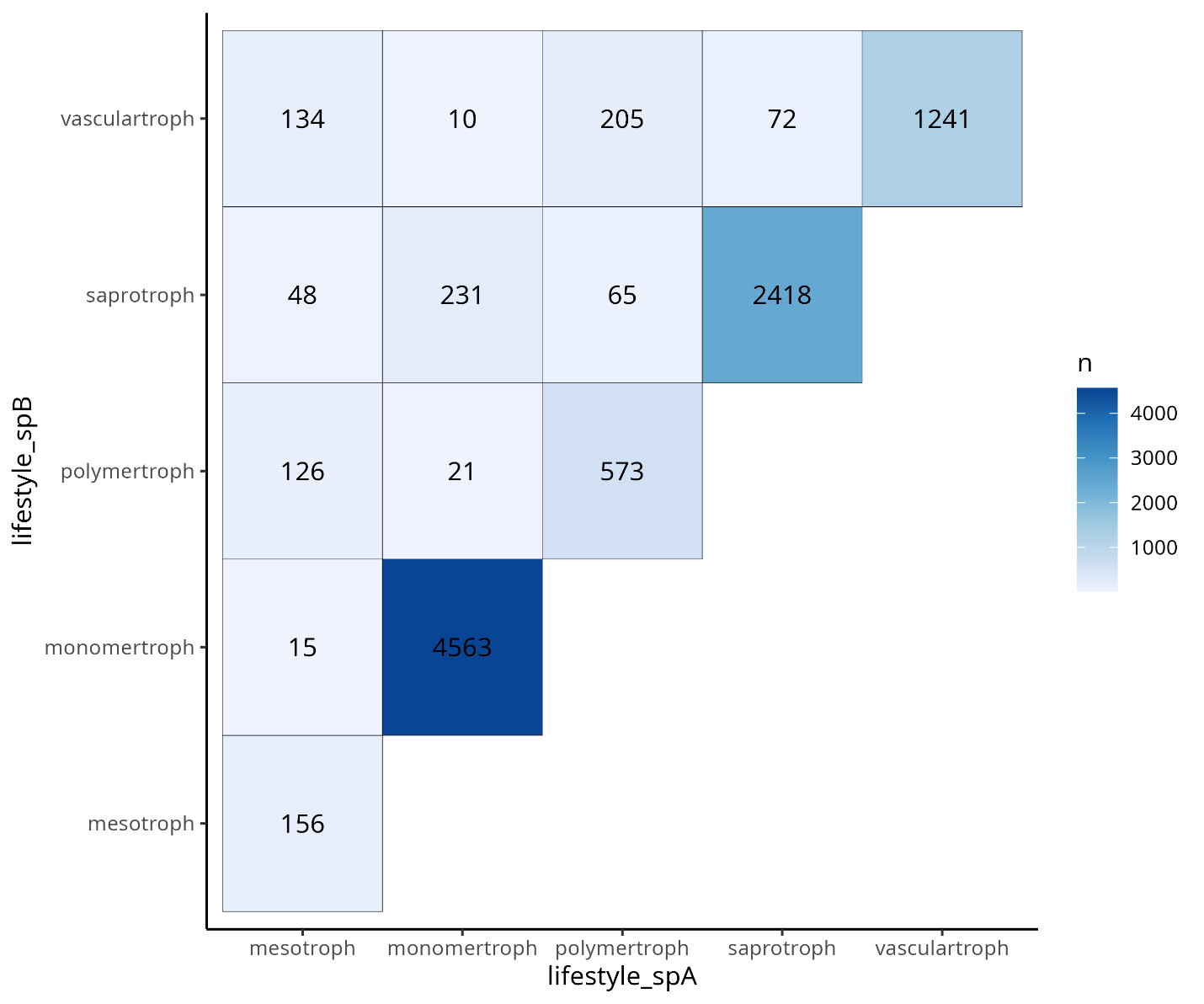

**Figure S4 – Tile plot displays the number of horizontally transferred transposable elements (HTTs) within and between different fungal lifestyles.** Number of HTTs is measured as the sum of number of hit communities observed between and within lifestyles (total number of hit communities = 11,001). Mesotrophs roughly translate to hemibiotrophs, monomertrophs to biotrophs, polymertrophs to necrotrophs and vasculartrophs to hemibiotrophs as well as wilts, rots and anthracnoses.

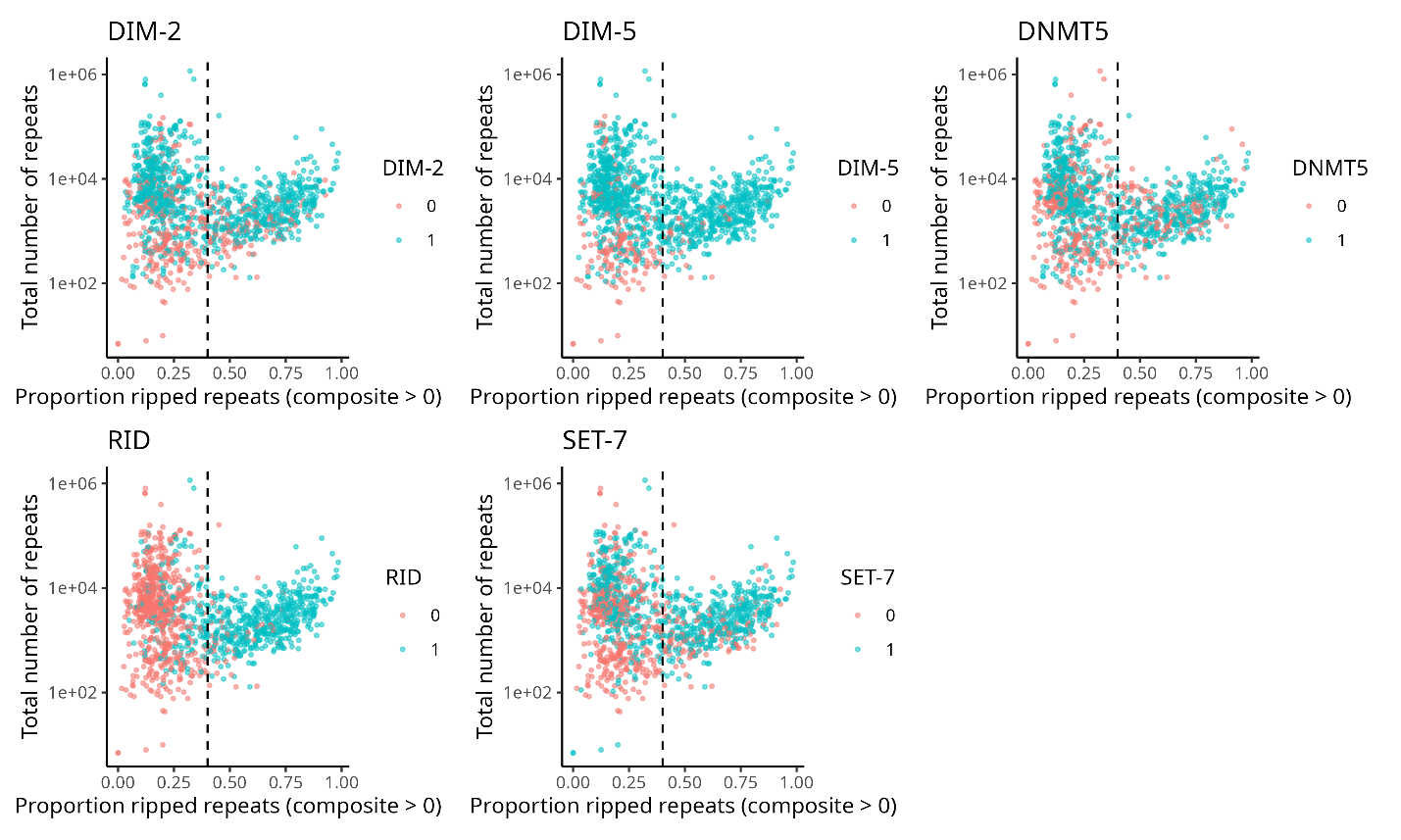

**Figure S5 – Presence and absence of five key enzymes involved in genome defenses.** Per protein and genome, the total number of repeats and proportion of ripped repeats is plotted. Note that here repeats include transposable elements (TEs), low complexity regions, RNAs, and G/A/GA-rich regions. Ripping of repeats was measured based on the composite index per repeat, with composite indices greater than 1 indicating RIP-signature^4^. We assessed correlations of the presence/absence of proteins with proportion of ripped repeats using Wilcoxon rank sum tests (DIM-2; p = 3.0e-4, DIM-5; p = 3.0e-16, DNMT5; p =3.1e-10, RID; p = 2.5e-150, SET-7; p = 3.3e-29). Abbreviations: DIM-2; DNA (cytosine-5-)-methyltransferase, DIM-5; Histone-lysine methyltransferase, DNMT5; DNA (cytosine-5-)-methyltransferase, RID; putative cytosine methyltransferase (RID stands for RIp Defective), SET-7; set-domain histone methyltransferase-7.

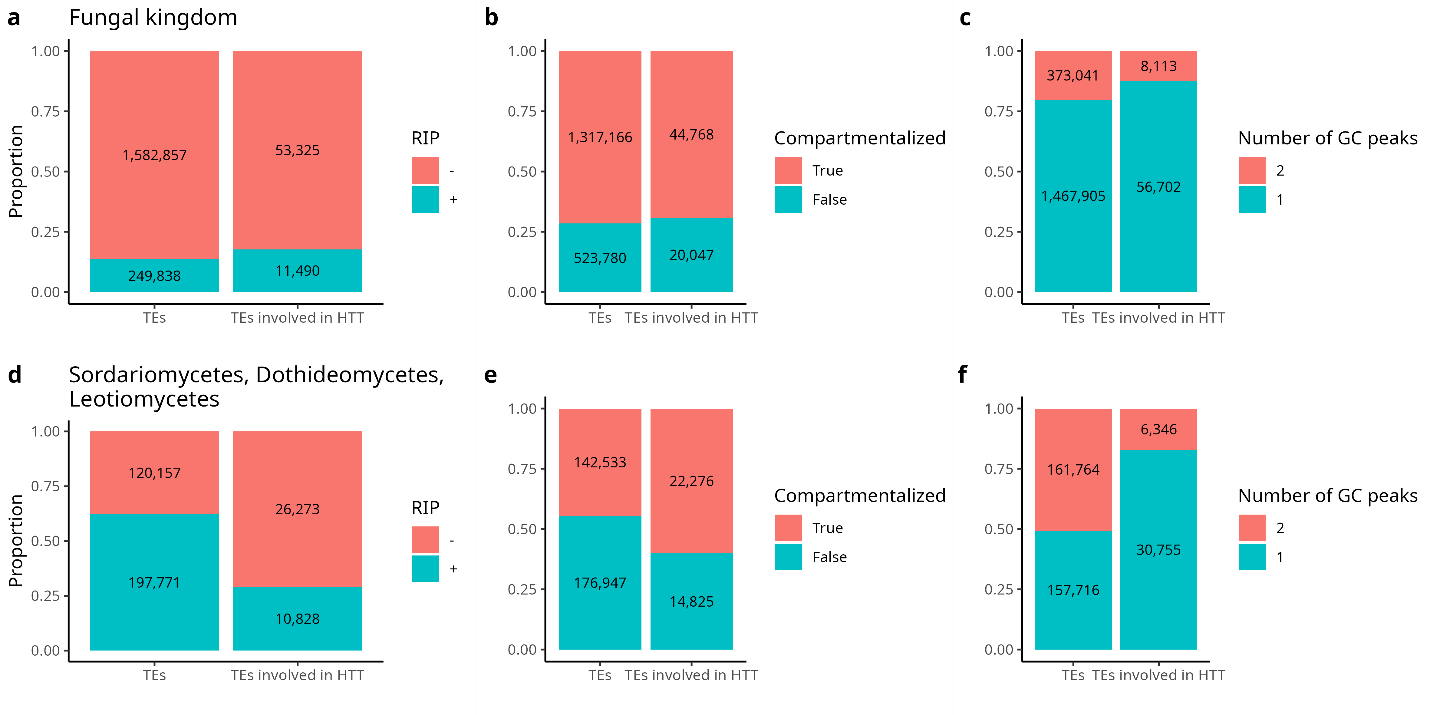

**Figure S6 - Enrichment analyses for different genome defenses and genome organizations.** Enrichment analyses were performed for RIP-proficient and RIP-deficient genomes (**a**,**d**) and for compartmentalized genomes (**b**,**e**), on the level of the fungal kingdom (**a,b**) and for fungal lineages enriched in transposable elements (TEs) involved in horizontal transfer of transposable elements (HTT) (**d,e**, see **Fig. 4a**). Note that too recently diverged genomes were collapsed and treated as a single genome, and some collapsed nodes have been removed from the analysis due to having inconsistent genome defenses or compartmentalization. The number of transposable elements (All TEs; TEs considered in the detection of HTT, n = 1,832,695 for fungal kingdom and n = 317,928 for Sordariomycetes, Dothideomycetes and Leotiomycetes, **Fig. S1**) as well as the number of TEs involved in HTTs (n = 64,815 for fungal kingdom and n = 37,101 for Sordariomycetes, Dothideomycetes and Leotiomycetes) are denoted in the plots. We performed enrichment tests (one-sided Fisher’s exact tests) with Bonferroni-Hochberg multiple testing correction per genomic feature for both the kingdom level as for the Sordariomycetes, Dothideomycetes and Leotiomycetes.

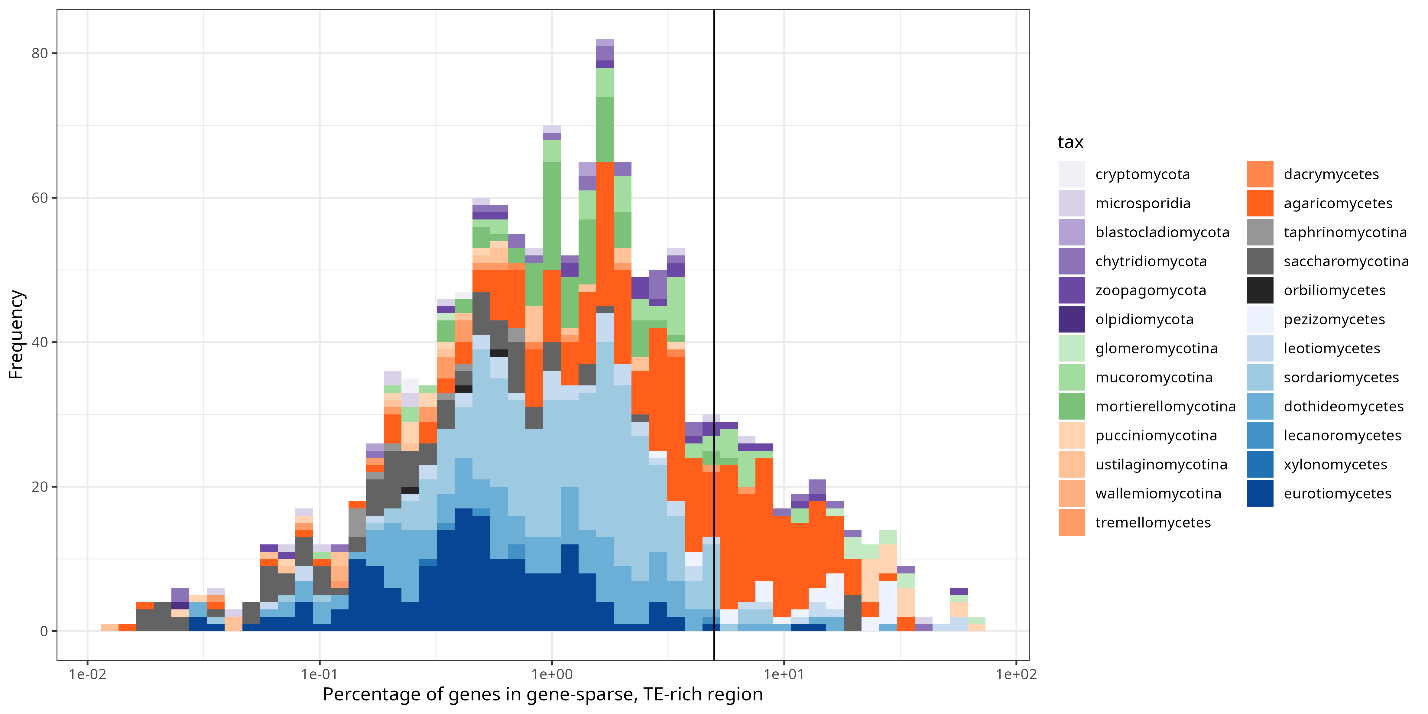

**Figure S7 – Number and taxonomic distribution of compartmentalized genomes.** Percentage of genes in gene-sparse and TE-rich region plotted per genome, with the frequency shown on the y-axis, colored by fungal lineage. If more than 5% of genes are located in gene-sparse and TE-rich regions, i.e. a neighboring gene is localized >5Kb and the closest TE is within 1Kb, the genomes are considered compartmentalized; 5% threshold is indicated with a vertical line.

**References**

1.	Clauset, A., Newman, M. E. J. & Moore, C. Finding community structure in very large networks. *Phys. Rev. E* **70**, 066111 (2004).

2.	Csardi, G. & Nepusz, T. The igraph software package for complex network research. *Available Igraphorg* **Complex Systems**, 1695 (2005).

3.	Traag, V. A., Waltman, L. & van Eck, N. J. From Louvain to Leiden: guaranteeing well-connected communities. *Sci. Rep.* **9**, 5233 (2019).

4.	van Wyk, S. *et al.* The RIPper, a web-based tool for genome-wide quantification of Repeat-Induced Point (RIP) mutations. *PeerJ* **7**, e7447 (2019).
